# Elevated genetic load is associated with reduced bite force in insular lizard populations

**DOI:** 10.64898/2026.10.07.752543

**Authors:** Lisa Van Linden, Ioanna Gavriilidi, Francisco Campuzano Jiménez, Zoë Maesen, Anamaria Štambuk, Anthony Herrel, Raoul Van Damme, Hannes Svardal

## Abstract

Demographic history can profoundly shape genomic variation by altering levels of inbreeding, the strength of genetic drift, and the efficacy of purifying selection, thereby influencing both the accumulation and expression of deleterious variation. Yet the extent to which these genomic consequences translate into organismal performance remains poorly understood. Here, we investigated how contrasting demographic histories shape genomic diversity, inbreeding, genetic load, and performance across mainland and insular populations of the Italian wall lizard, *Podarcis siculus*. Using whole-genome sequence data from 263 wild-caught individuals and field-based performance measurements from 134 lizards, we characterised population structure and demographic history, quantified genomic diversity, runs of homozygosity, and deleterious variation, and tested associations between genomic inbreeding and performance traits. Island populations showed reduced genetic diversity, increased autozygosity, and elevated genetic load relative to mainland populations, consistent with stronger genetic drift and reduced efficacy of purifying selection. Patterns of deleterious variation were consistent with purging of strongly, but not mildly, deleterious variants. Greater genomic autozygosity was associated with reduced bite force in island lizards, a pattern consistent with inbreeding depression, whereas sprint speed, sperm quality, immunocompetence, and spatial cognition showed no significant association with inbreeding. Genome-wide mapping of runs of homozygosity further identified multiple genomic regions associated with reduced bite performance. Together, these results show how demographic history shapes genomic diversity, homozygosity, and deleterious variation in wild vertebrate populations, while revealing that phenotypic consequences can be trait-specific, a pattern that may reflect differences in genetic architecture and the extent of environmental influence on trait variation.

## Introduction

Population genetic theory predicts that island populations exhibit elevated genetic load and reduced mean fitness relative to mainland populations (Frankham, 1998; Keller & Waller, 2002). Small founding populations and subsequent demographic bottlenecks reduce effective population size, increasing genetic drift and reducing the efficacy of purifying selection, thereby promoting the accumulation of deleterious variation (drift load; Allendorf et al., 2022; Whitlock & Davis, 2011). In addition, small population size increases inbreeding (Crow, 1970; Soulé, 1987), leading to loss of heterozygosity and the expression of recessive deleterious alleles (inbreeding depression; Charlesworth & Willis, 2009). If selection is insufficient to purge deleterious variants (Dussex et al., 2023; Hedrick & Garcia-Dorado, 2016), these processes are expected to reduce fitness and increase extinction risk in island populations (Hedrick & Kalinowski, 2000; Kardos et al., 2021; Tanaka, 1997).

Empirical evidence for these predictions is mixed. While many studies report reduced genetic diversity, increased inbreeding, and elevated estimates of genetic load in island populations compared to mainland counterparts (e.g. Cavill et al., 2024; Dussex et al., 2021; Lavanchy, Cumer, et al., 2024; Robinson et al., 2018; Van Noordwijk & Scharloo, 1981), others find weak, context-dependent (e.g. Gabrielli et al., 2026; García-Verdugo et al., 2015; James et al., 2016a; Machado et al., 2022; Otalora et al., 2024; Vlček et al., 2025), or even contrasting patterns (e.g. Michaelides et al., 2025; Tsumura & Ohba, 1993). This variation suggests that the genetic consequences of isolation depend on a variety of factors, such as colonisation history, demographic trajectories, and the extent of purging, yet these processes remain insufficiently resolved in most natural systems (reviewed in Gabrielli et al., 2026).

Recent advances in population genomics now allow demographic history, genomic inbreeding, and deleterious variation to be characterised jointly in wild populations using genome-wide data, including through identity-by-descent segments (Kardos et al., 2015). Combining these genomic approaches with measurements of organismal traits provides an opportunity to test how demographic history and associated genomic erosion translate into variation in phenotypic performance in comparative mainland-island systems.

However, how increased inbreeding and genetic load are reflected in organismal performance remains poorly understood. Most studies focus either on morphological indicators of genetic health (e.g. fluctuating asymmetry, morphological aberrations; Eldridge et al., 1999; Plećaš et al., 2024; Robinson et al., 2018) or on high-level fitness components such as survival and reproduction (e.g. Brewer et al., 1990; Hasselgren et al., 2024). Measures of whole-organism performance provide an important intermediate level linking variation in condition and functional capacity to ecologically relevant fitness components (Arnold, 1983), yet such integrative comparisons remain rare across insular and mainland populations.

Here, we address this knowledge gap using the Italian wall lizard (*Podarcis siculus*) as a model system. Owing to its widespread distribution across mainland Italy and numerous Mediterranean islands (Podnar et al., 2005; Senczuk et al., 2017), populations vary substantially in demographic history, genetic diversity, and levels of isolation (Sherpa, Salvi, et al., 2023). In a mainland-island comparative framework, we reconstruct population structure and demographic history, quantify genomic diversity and load, and estimate individual inbreeding. We then test how these patterns relate to variation in multiple performance traits and characterise the architecture of inbreeding depression in the genome. This integrated approach allows us to assess whether differences in demographic history correspond to differences in genomic load and whether those are associated with variation in organismal performance.

## Results

Whole-genome sequencing data were generated for 205 *P. siculus* individuals and combined with 58 previously published genomes from Van Linden et al. (2025), resulted in a final dataset of 263 samples from thirteen populations (Figure 1A). The median sequencing coverage was 15.6×, with an average of 99.55% of reads mapping to the reference genome (Table S2). Following variant calling and filtering, ∼130 million SNPs were retained for downstream genomic analyses.

**Figure 1:**
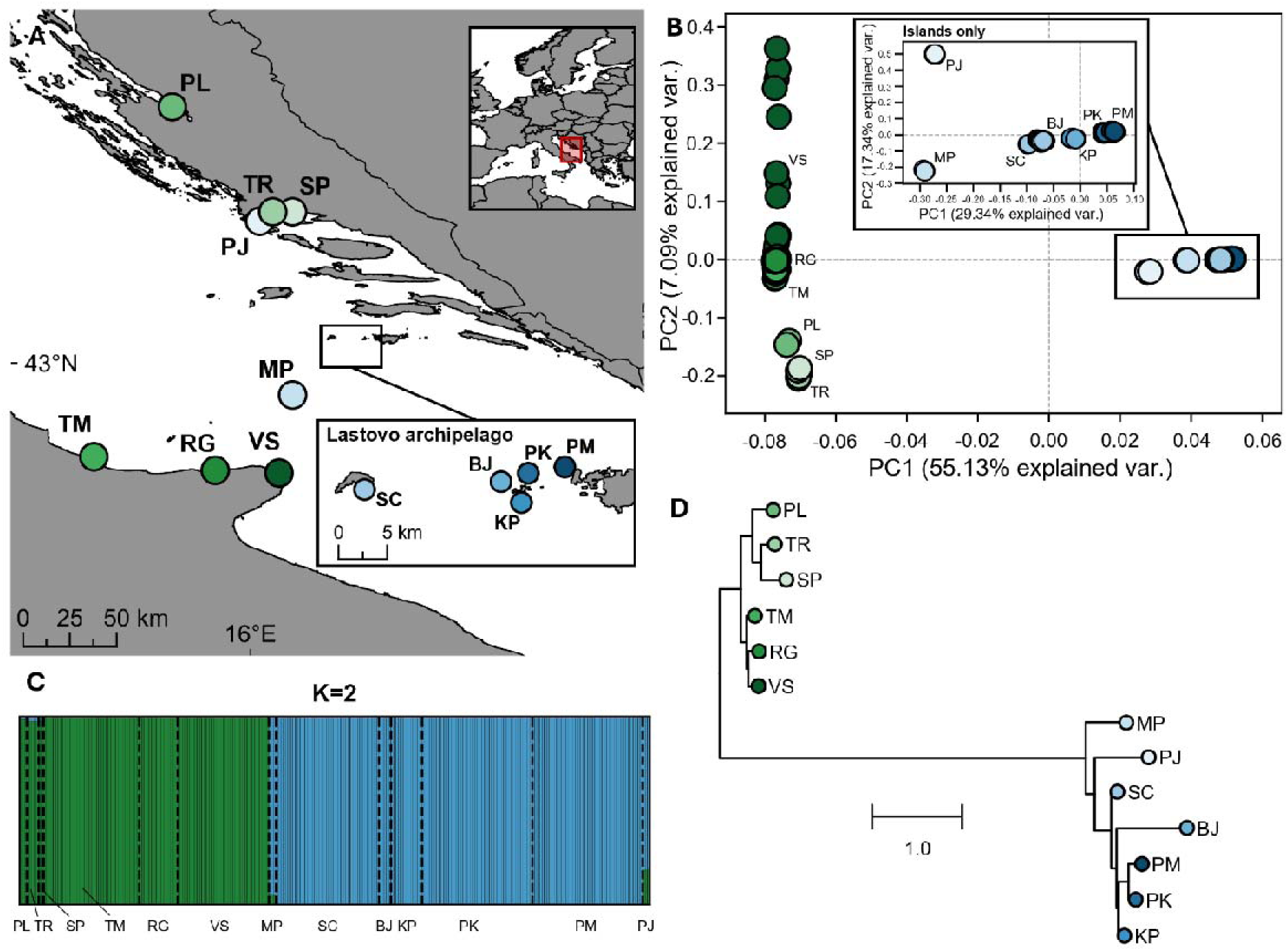
Sampling and population structure. **A.** Map of the sampling locations of the mainland (green shades) and island (blue shades) *Podarcis siculus* populations included in this study. The inset on the top right provides the broader geographical context of the study area (red square). A more detailed view of the sampled island populations in the Lastovo archipelago is shown at the lower right. See Supplementary Information for detailed coordinates of the sampling locations, sampling years, and sample sizes for all populations. **B.** Principal component analysis (PCA) for all *P. siculus* samples, with an inset plot showing the PCA of only the island populations. Each dot represents an individual. PC axis labels show % variance explained. **C.** ADMIXTURE plot with K=2 (lowest cross-validation error). Each bar represents a single *P. siculus* individual, with dashed lines marking population boundaries. Mainland and island genetic clusters are coloured in green and blue, respectively. **D.** Phylogenetic tree showing the genetic relationships among the sampled populations, derived from the consensus tree constructed with ASTRAL-III from 13705 maximum likelihood trees of 100KB genomic windows, with *P. muralis* as the outgroup (not shown). The scale bar represents coalescent units.

### Population structure and phylogenetic relationships

We first characterised genome-wide population structure and genetic relationships among the sampled mainland and island populations. The genomic PCA revealed strong divergence between mainland and island populations of *P. siculus*, loading on PC1 and accounting for 55.13% of the observed genetic variation (Figure 1B). PC2 (7.09% variation explained) primarily captured differentiation between Croatian and Italian mainland populations and relatively high within-population variation in the Vieste (VS; Italy) population (Figure 1B). Among island populations, genetic structure largely mirrored geographic proximity, with each island forming a distinct cluster (PC1=23.34%, PC2=17.34% variation explained). Within the Lastovo archipelago, individuals from Pod Kopište (PK) and Pod Mrčaru (PM) clustered closely together, consistent with their shared genetic history following the documented translocation of 10 individuals from PK to PM (Nevo et al., 1972).

Model-based clustering with ADMIXTURE analysis supported this structure, identifying two main genetic clusters (K=2; lowest CV error, Table S3) separating mainland and island populations (Figure 1C). Island populations closest to the mainland (i.e. Pijavica, PJ; Mala Palagruža, MP) showed mixed ancestry, consistent with (limited) gene flow involving both Italian and Croatian mainland populations, as also supported by D-statistics (Table S4). For runs with higher numbers of genetic clusters, Italian mainland populations exhibited substantial within-population variation and were clearly separated from Croatian mainland populations (Figure S1). Among the islands, genetic structure likely reflects the combined effects of genetic isolation and independent drift, with several populations forming distinct ancestry components at higher values of K and showing limited evidence of recent admixture in D-statistics (Figure S1; Table S4).

Genome-wide genetic differentiation further quantified the extent of divergence between mainland and island populations. Genome-wide differentiation was high between mainland and island populations (mean F_st_=0.42), whereas divergence between Italian and Croatian mainland populations was low (F_st_=0.040). Among island groups (MP, PJ, and the Lastovo archipelago), pairwise F_st_ ranged from 0.23 (MP vs. Lastovo archipelago) to 0.32 (PJ vs. other island populations).

These patterns are mirrored in the phylogeny (Figure 1D), which clearly separates mainland and island populations and shows island populations as distinct lineages. Relationships within both mainland and island clusters broadly reflect geographic proximity. The ancestral branch leading to the island populations is relatively long, consistent with strong genetic drift.

### Demographic histories of mainland and island populations

We next reconstructed historical effective population size (N_e_) trajectories to assess whether differences in genetic diversity between mainland and island populations reflected contrasting demographic histories. Population history inferred with PHLASH showed broadly similar effective population size (N_e_) trajectories across all populations in deep time (>2 MYA), consistent with shared ancestral dynamics (Figure 2). The earliest major divergence in effective population sizes between mainland and island lineages occurs during the Mid-Pleistocene Transition (MPT; ∼0.7-1.25 MYA). During this period, N_e_ increased sharply in the mainland populations, particularly in the Italian mainland populations, where it exceeded 10 million, whereas the Croatian mainland populations showed a more moderate increase. Within the mainland clade, the Croatian populations showed consistently lower N_e_ through time after the Penultimate Glacial Maximum (PGM; ∼135-194 KYA).

**Figure 2:**
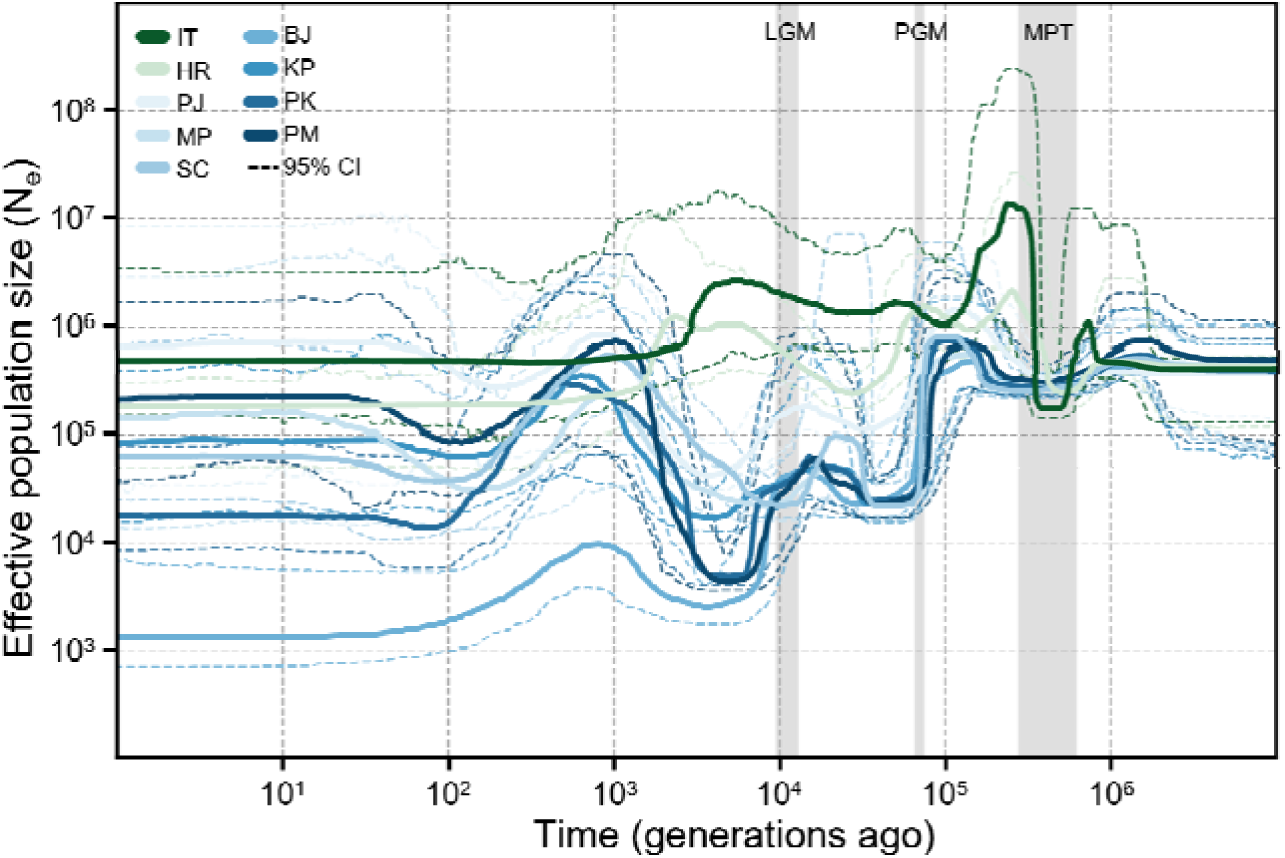
Demographic history. Temporal changes in effective population size (N_e_) inferred with PHLASH for mainland (green) and island (blue) populations. Time is shown in generations before present, assuming a generation time of two years. Dashed lines represent 95% credible intervals. Abbreviations: LGM: Last Glacial Maximum; PGM: Penultimate Glacial Maximum; MPT: Mid-Pleistocene Transition.

Island populations showed a moderate increase in N_e_ between 400 and 200 KYA, followed by a decline during the PGM to ∼25.000. Trajectories remained broadly similar across island populations until the Last Glacial Maximum (LGM; ∼20-26 KYA), after which they appear to diverge, although 95% credible intervals were wide and largely overlapping. Post-LGM trajectories generally indicate population declines followed by recovery, although the timing and magnitude of these changes vary among populations. Recent demographic inference (<100 years) remains poorly resolved, as is evident from the trajectories’ substantial uncertainty and largely overlapping credible intervals across populations, and the absence of a clear demographic split between the PK and PM populations despite their known recent divergence (i.e. 1971, Nevo et al. 1972). Restricted sample sizes in our dataset likely constrains inference of recent coalescent events, resulting in wide credible intervals and reduced resolution toward the present.

### Genomic diversity, inbreeding, and genetic load

We compared genome-wide genetic diversity and genomic inbreeding between mainland and island populations, and examined their relationship across individuals. Island populations of *P. siculus* showed substantially lower genome-wide heterozygosity than mainland populations (0.0019 ± 0.000024 vs. 0.0067 ± 0.00024; LMER, t=20.25, p<0.001; Figures 3A, S2). Consistent with this, island individuals exhibited higher proportion of their genome in ROHs, with a mean F_ROH_ of 0.59 ± 0.0047 compared to 0.11 ± 0.0033 in mainland individuals (GLMER, z=-27.49, p<0.001; Figure 3B). Genome-wide heterozygosity declined with increasing F_ROH_ (LMER, t=-20.58, p<0.001), with a significantly stronger negative relationship in mainland than in island populations (LMER interaction, t=-12.61, p<0.001).

**Figure 3:**
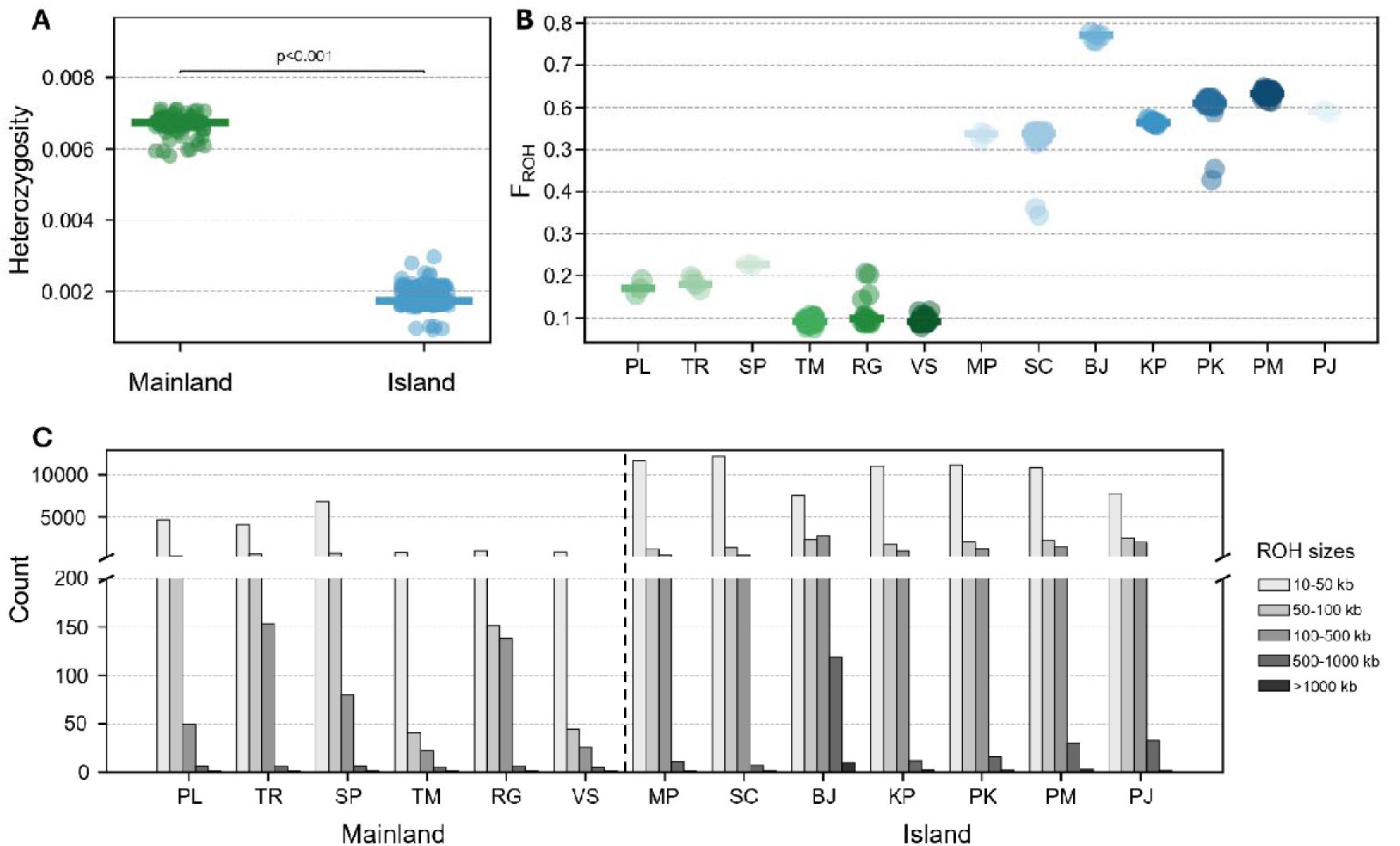
Genomic diversity and inbreeding levels. **A.** Mean genome-wide heterozygosity of mainland (green) and island (blue) *Podarcis siculus* individuals, estimated with ANGSD v0.941 from genotype likelihoods and folded site frequency spectra computed in 100 kb non-overlapping windows. Points show individual mean heterozygosity (averaged across genome windows). Horizontal lines represent mainland and island median heterozygosity. **B.** Individual F_ROH_ estimates for *P. siculus* individuals from mainland (green shades) and island (blue shades) populations from runs-of-homozygosity (ROHs) inferred with BCFtools/RoH from genotypes likelihoods. Horizontal coloured lines represent population median F_ROH_. **C.** Mean number of ROHs per population across size classes (10-50 kb, 50-100 kb, 100-500 kb, 500-1000 kb, and >1000 kb) found in the genomes of the different island and mainland populations of *P. siculus*, inferred with BCFtools/RoH. Bars show population averages, with darker shades indicating larger ROH size classes. The dashed vertical line separates mainland (left) and island (right) populations.

We then examined the number, length, and genomic distribution of ROHs to determine whether the higher inbreeding of island populations reflected differences in the timing and genomic distribution of inbreeding. Island populations harboured more ROHs across all size classes than mainland populations (Figure 3C; all p<0.001). Genome-wide distributions of ROH were largely distinct between mainland and island populations (pairwise Pearson’s correlation r=0.00). Mainland populations showed weak similarity in ROH landscape (mean r=0.14), whereas island populations exhibited higher similarity in ROH regions (mean r=0.37, up to 0.85), with ∼2% of the genome fixed in ROH across islands (Figure S3).The distribution of the shortest ROH classes (10-50 kb and 50-100 kb) indicated that differences in inbreeding levels between Italian mainland and island populations primarily reflect demographic processes occurring ∼400-1000 generations ago, rather than recent inbreeding in the last few generations, as differences among populations decrease with increasing ROH length. Consistent with this pattern, population-level inbreeding coefficients (F), estimated from deviations from Hardy-Weinberg equilibrium, were close to zero across populations (Table S5). Among mainland populations, Croatian populations exhibited higher proportions of ROHs than Italian populations (Figure 3B), with numbers of shorter ROHs (50-100 kb) comparable to those observed in island populations (Figure 3C).

Among island populations, lizards from BJ showed the lowest heterozygosity (mean= 0.00098) (Figure S2) and the highest inbreeding levels, with a mean F_ROH_ of 0.77 and the greatest number of ROHs across all length classes (Figure 3B, C). ROHs of 500-1000 kb and >1000 kb, indicative of relatively recent inbreeding within the last ∼30 generations, were particularly abundant in this population. The PM population showed the second-highest inbreeding levels, although heterozygosity and ROH patterns were broadly comparable to those of the source population PK and other island populations from the Lastovo archipelago (Figures 3B, 3C, S2).

We next assessed whether the reduced genetic diversity and increased genomic inbreeding of island populations were accompanied by elevated genetic load. Island lizards exhibited higher genetic load than mainland lizards (Figure 4). Both masked (heterozygous) and realised (homozygous) load were elevated in island individuals, as reflected by higher DEL/SYN and LOF/SYN ratios (Figure 4A; all p<0.05). Consistent with this pattern, island populations showed a shift in allele frequency distributions indicative of increased load. The ratio of nonsynonymous to synonymous variants (NS/S) was higher across allele frequency bins, and deleterious variants (DEL and LOF) were more often found at higher frequencies compared to mainland populations (Figure 4B).

**Figure 4:**
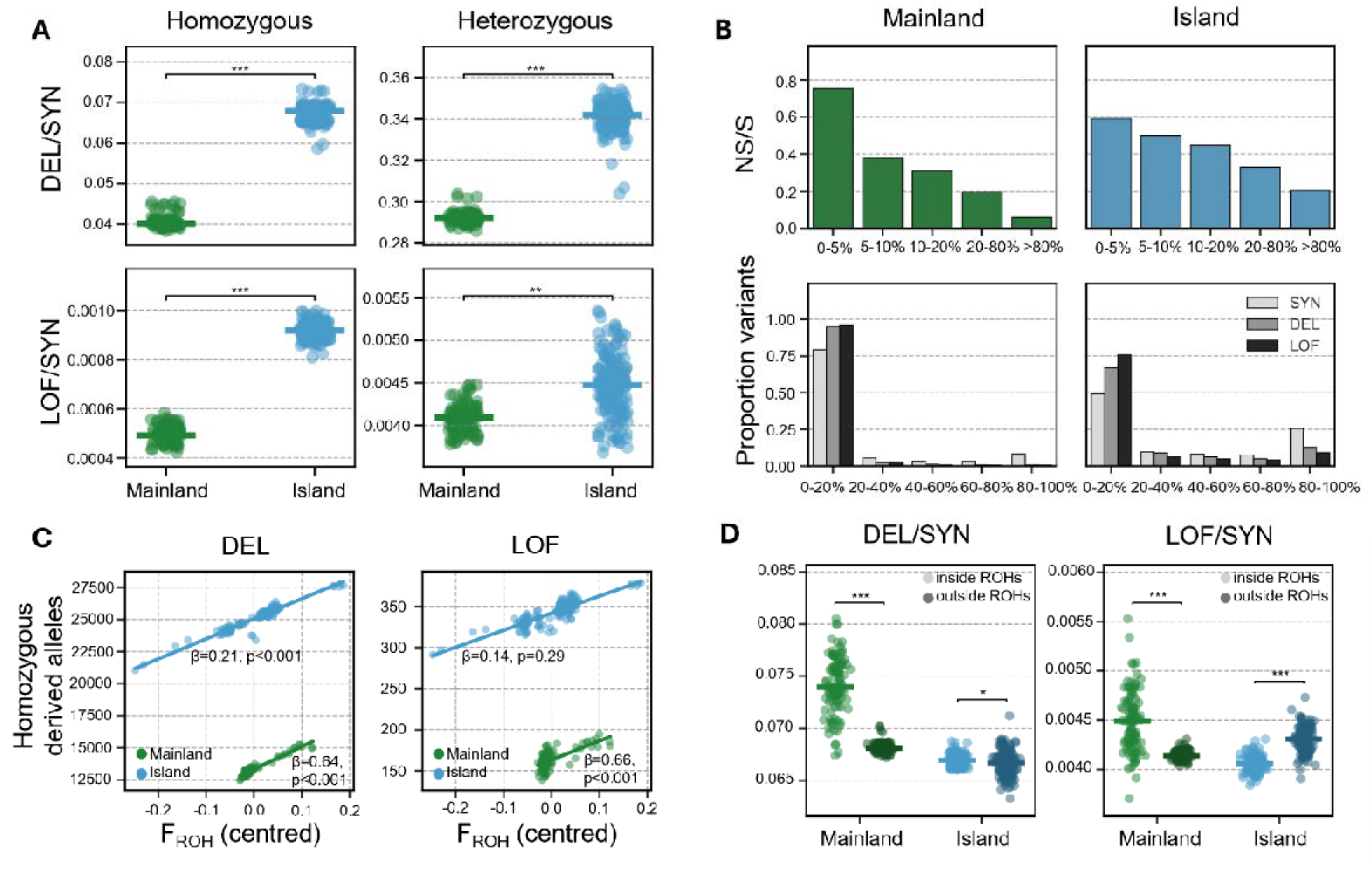
Genetic load. **A.** DEL/SYN and LOF/SYN ratios for homozygous (realised) and heterozygous (masked) variants in mainland (green) and island (blue) *Podarcis siculus* individuals. Variants were classified using SnpEff into synonymous (SYN), missense (DEL), and loss-of-function (LOF) categories. Points represent individuals, and horizontal lines indicate group medians. **B.** Allele frequency distributions for mainland and island populations based on the unfolded site frequency spectrum (SFS). Top panels show nonsynonymous-to-synonymous (NS/S) ratios across derived allele frequency bins, and bottom panels show the proportion of SYN, DEL, and LOF variants within each bin. **C.** Relationship between genomic inbreeding (F_ROH_, centred) and the number of homozygous derived alleles for DEL and LOF variants in mainland and island individuals. Points represent individuals, and lines show linear mixed-effects model fits, natural log-transformed, controlling for synonymous variation and including population as a random effect. **D.** DEL/SYN and LOF/SYN ratios within and outside ROHs for mainland and island populations. Ratios were calculated separately for genomic regions inside and outside ROHs, with allele counts weighted additively (heterozygotes=1, homozygotes=2). Points represent individuals, and horizontal lines indicate group medians; lighter shades denote values inside ROHs and darker shades denote values outside ROHs.

We further tested whether individual variation in genomic inbreeding was associated with the accumulation of deleterious variants. Homozygous deleterious (DEL) load increased with genomic inbreeding (F_ROH_) in both mainland and island populations (LMER; mainland: β=0.64, t=10.90, p<0.001; island: β=0.21, t=6.43, p<0.001; Figure 4C). In contrast, for LOF variants, the relationship with F_ROH_ was weak and not significant in island populations (LMER: β=0.14, t=1.06, p=0.29), but significantly stronger in mainland populations (LMER: β=0.66; t=3.75, p<0.001).

Finally, we compared the distribution of deleterious variation inside and outside ROHs to assess whether homozygosity was associated with the exposure of genetic load. The distribution of genetic load differed between ROH and non-ROH regions, variant types, and mainland and island populations (Figure 4D). For deleterious variants, DEL/SYN ratios were higher inside ROHs (LMER: t=-2.22, p<0.05), with this enrichment being more pronounced in mainland populations (LMER: t=-22.96, p<0.001). In contrast, patterns for loss-of-function variants differed between regions: LOF/SYN ratios were higher outside ROHs in island populations (LMER: t=14.16, p<0.05), whereas the opposite pattern was observed in mainland populations, i.e. LOF/SYN ratios were elevated within ROHs (LMER: t=-20.98, p<0.05).

### Associations between genomic inbreeding and lizard performance

We tested whether the elevated genomic inbreeding observed in island populations was associated with reduced performance across traits linked to fitness. In island lizards, greater genomic inbreeding (F_ROH_) was associated with a significant reduction in bite force (β=-3.55, 95% CI: -6.20 to -1.12, pMCMC = 0.003), consistent with inbreeding depression in bite performance. In contrast, no such relationship was detected in mainland lizards (β = 4.37, 95% CI: -3.69 to 12.65, pMCMC = 0.296) (Figure 5; Table S7A). This association in island lizards persisted after centring F_ROH_ within populations (β = -7.12, 95% CI: -14.14 to -0.35, pMCMC = 0.042), indicating that it was not solely driven by differences in mean F_ROH_ among island populations. The F_ROH_-bite force association was negative within each population, with F_ROH_ × population interactions not supported (Figure S4; Table S7).

**Figure 5:**
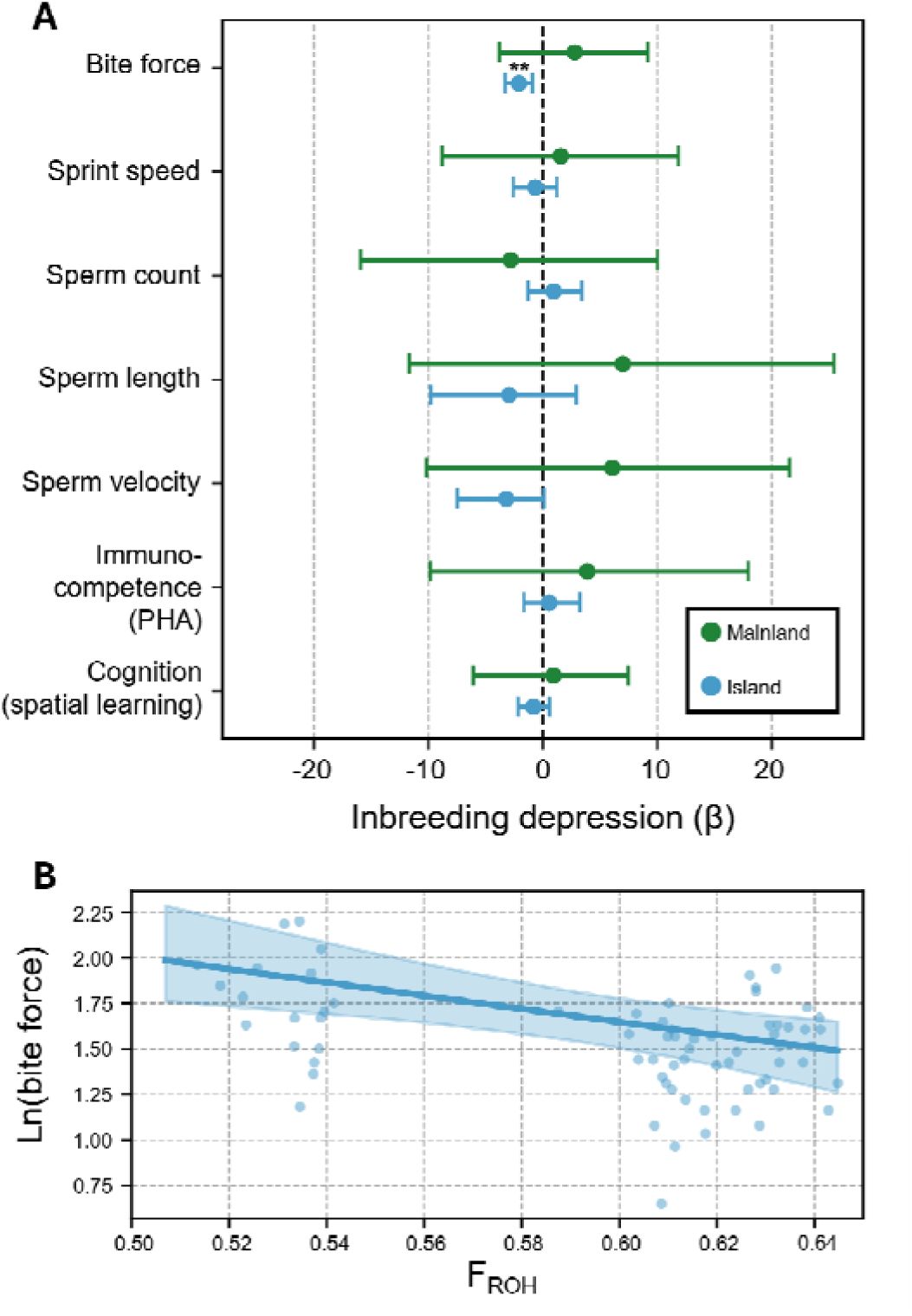
Inbreeding depression. **A.** Standardised inbreeding depression estimates (β ± 95% credible intervals) for biting and running performance, immune, reproductive, and cognitive traits in male *Podarcis siculus* using F_ROH_. Mainland populations are shown in green and island populations in blue; significant effects (credible intervals not overlapping zero) are indicated with asterisks. Full MCMCglmm model outputs are found in Table S7-13. **B.** Relationship between F_ROH_ and (natural log-transformed) bite force in island lizards. Points represent individuals; the line and shaded area show posterior mean predictions and 95% credible intervals from the MCMCglmm model, with head size held constant at its mean. Population-averaged trait values (F_ROH_ and performance) are found in Table S6. See Figure S4 for island population-specific relationships between F_ROH_ and bite force.

No evidence for inbreeding depression was found for other performance traits, including sprint speed, sperm quality, immunocompetence, and spatial cognition, in either island or mainland populations (N=134 individuals assessed for performance, all 95% CI overlapping zero and pMCMC > 0.05; Figure 5; Tables S7-13).

### Genomic basis of inbreeding depression in bite performance

Because genomic inbreeding was associated with reduced bite force in island lizards, we next tested whether specific genomic regions contributed to this pattern by examining genome-wide associations between ROHs and bite force in mainland and island male *P. siculus*. We detected significant associations in both mainland and island populations.

In mainland populations, we identified three significant windows (exceeding the adjusted Bonferroni threshold), all showing negative effects of ROH presence on bite force, indicating reduced performance when these regions were homozygous. These windows were located on chromosomes 3 and 10 (Figure 6A) and overlapped ROH in both the TM and VS populations (Figure S5).

**Figure 6:**
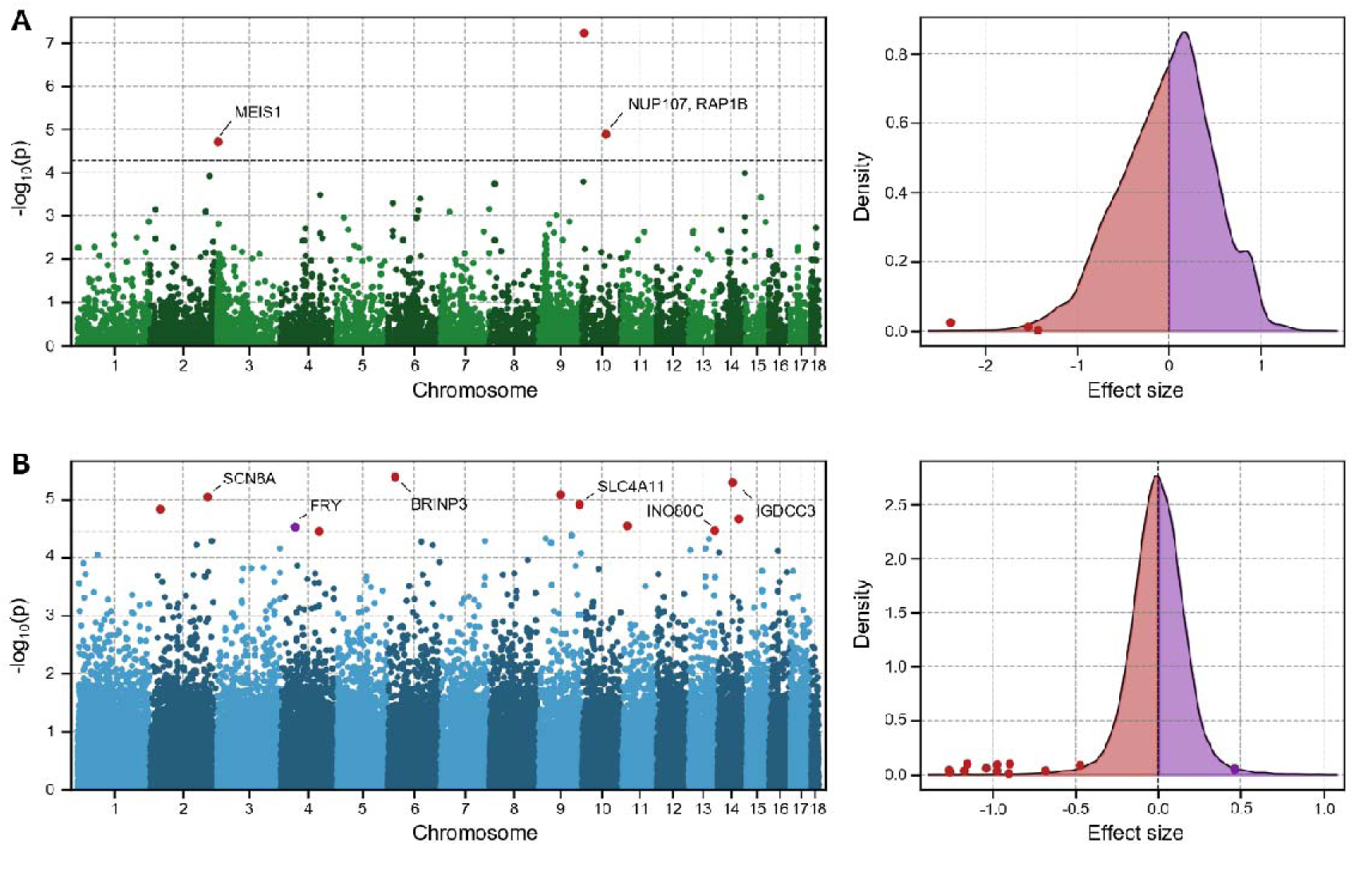
Genetic architecture of inbreeding depression in bite force. ROH-GWAS Manhattan plots (left) and effect size distributions (right) for **A.** mainland and **B.** island populations of *Podarcis siculus*. Manhattan plots show window-based ROH_status_ associations with bite force across the genome, with SNP windows exceeding the SimpleM-adjusted Bonferroni threshold (α/Neff; black dashed line) highlighted as points in red (negative effect) and purple (positive effect). Density plots show the genome-wide distribution of effect sizes, with significant windows indicated as points. Genes from the *P. muralis* to *P. siculus* Liftoff annotation overlapping significant windows are labelled (Table S14).

In island populations, thirteen significant windows were distributed across chromosomes 2, 4, 6, 9, 11, 13, and 14. Eleven of these showed negative associations between ROH overlap and bite force, consistent with inbreeding depression (Figure 6B), and were mostly fixed for ROHs in the PK and PM populations (Figure S5).

Genome-wide distributions of window-specific ROH effect sizes were centred near zero in both mainland (mean=-0.025, median=0.018) and island populations (mean=-0.0015, median=-0.00065), although island populations exhibited a more pronounced tail of negative effects (Figure 6). We identified four protein-coding genes overlapping significant windows in mainland populations (MEIS1, LOC114604855, NUP107, and RAP1B) and nine in island populations (LOC114588424, LOC114588406, SCN8A, FRY, BRINP3, SLC4A11, LOC114606797, INO80C, and IGDCC3; Figure 6, Table S14).

## Discussion

We investigated how mainland and insular demographic histories shape genome-wide diversity, inbreeding, genetic load, and inbreeding depression across wild *P. siculus* populations.

### Distinct demographic trajectories shape differences in genetic diversity and load between mainland and island lizards

Population genetic analyses revealed strong differentiation between insular and mainland *P. siculus*, with genetic clustering, phylogenetic relationships, and admixture patterns largely reflecting geographic proximity and insular isolation, typical for Mediterranean *Podarcis* populations (e.g. Giovannotti et al., 2010; Hurston et al., 2009; Podnar et al., 2004, 2005; Runemark et al., 2012; Sabolić, 2021; Senczuk et al., 2017; Sherpa, Paris, et al., 2023; Sherpa, Salvi, et al., 2023; Van Linden et al., 2025) (Figures 1, S1; Table S4). Effective population size trajectories suggest that mainland and island lineages diverged during the Mid-Pleistocene Transition (∼0.7-1.25 MYA) (Figure 2), consistent with previous estimates (Senczuk et al., 2017; Sherpa, Salvi, et al., 2023). This period of climatic oscillations likely promoted lineage divergence through repeated isolation in Mediterranean refugia (Böhme et al., 2007; Canestrelli et al., 2012; Salvi et al., 2013; Senczuk et al., 2017). Subsequently, island populations experienced repeated contractions and expansions, likely reflecting climatic oscillations and island founder events with demographic instability, whereas mainland populations maintained comparatively larger and more stable effective population sizes.

These contrasting demographic histories are expected to increase genetic drift and reduce the efficacy of purifying selection in island populations (Crow, 1970; Keller & Waller, 2002; Whitlock & Davis, 2011), leading to lower diversity, greater homozygosity, and higher genetic load relative to their mainland counterparts (Bortoluzzi et al., 2020; Chen et al., 2025; Lozier et al., 2023; Machado et al., 2022; Martin et al., 2023). Consistent with these predictions, island lizards had less than half the genome-wide heterozygosity of mainland individuals, up to 40% more and longer ROHs, and higher masked and realised genetic load (Figures 3, 4, S2). Population-level variation also broadly reflected demographic history: Croatian mainland lizards had higher F_ROH_ than Italian mainland lizards, in line with their lower long-term effective population sizes; lizards on the island of Bijelac, which had the smallest inferred effective population size, showed the greatest proportion of their genome in ROHs (Figures 2, 3, S2). Interestingly, a large portion of the ROH landscape was similar across all island populations, indicating shared drift dynamics (Figure S3).

Pod Mrčaru provides an informative contrast. Despite being founded by only ten individuals following its introduction in 1972 (Nevo et al., 1972), the population shows detectable but relatively modest genomic signatures of increased inbreeding compared with its source population, Pod Kopište, with only slightly reduced heterozygosity (consistent with previous assessments utilising reduced representation genomic data; Sabolić et al., 2024; Sherpa, Paris, et al., 2023) and slightly elevated F_ROH_ and ROH lengths (Figures 3, S2). This pattern is most consistent with a strong but short-lived founding bottleneck followed by rapid population expansion, potentially facilitated by the high reproductive output characteristic of *Podarcis* (Biaggini & Corti, 2019; Carretero, 2006; Ji & Braña, 2000; Oppliger et al., 2007; Rotger et al., 2023), which would have limited the loss of genetic diversity and accumulation of homozygosity. Because Pod Mrčaru is smaller than Pod Kopište, its present-day effective population size may nevertheless remain lower, in which case some continued loss of heterozygosity through drift could still occur (drift debt; Gargiulo et al., 2024). However, current effective population sizes are not known, and census size may be only weakly related to N_e_, so the extent of any remaining drift debt is uncertain.

Together, these results indicate that demographic history strongly shapes genomic diversity and inbreeding across *P. siculus*, while highlighting that the genomic consequences of population bottlenecks depend not only on the magnitude of population decline, but also on its duration and subsequent recovery. Short bottlenecks involving very small population sizes may leave different and more limited signatures than prolonged periods of small effective population size, particularly when followed by rapid population growth. Bottleneck severity alone therefore provides an incomplete indication of the extent and nature of genomic erosion experienced by a population. However, although rapid demographic recovery may limit the accumulation of long ROHs, it may not prevent the accumulation and expression of deleterious variation in island populations (Bertorelle et al., 2022; Frankham, 1998; Tanaka, 1997).

### Differential signatures of selection on deleterious variation

Insular *P. siculus* exhibited higher genetic load than mainland populations, with elevated DEL/SYN and LOF/SYN ratios and higher frequencies of putatively deleterious alleles (Figure 4A, B). These patterns are consistent with stronger genetic drift under small effective population sizes, which reduces the efficacy of purifying selection and promotes the accumulation of deleterious variants (Keller & Waller, 2002; Whitlock & Davis, 2011). The increase in both realised (i.e. homozygous) and masked (heterozygous) load further indicates a greater current fitness burden and a greater reservoir of recessive or partially recessive deleterious alleles that may become expressed under future inbreeding (Bertorelle et al., 2022; Dussex et al., 2023).

We found patterns consistent with some removal of strongly deleterious variation (i.e. purging) in island populations. In particular, LOF variants were relatively depleted within ROHs in island lizards, suggesting that strongly deleterious recessive alleles may have been preferentially removed from genomic regions that experienced repeated or persistent homozygosity during past population bottlenecks (Chen et al., 2025; Lan et al., 2025; Taylor et al., 2024; L. Yang et al., 2025). However, homozygous LOF load showed no clear relationship with contemporary genomic inbreeding (F_ROH_). We interpret this latter result cautiously, because variation in F_ROH_ among present-day individuals and populations reflects multiple aspects of demographic history and does not directly measure the historical exposure of deleterious alleles to selection. In particular, in small populations, drift may simultaneously increase homozygosity and the frequency of deleterious alleles, potentially obscuring signatures of purging.

Patterns for more weakly deleterious variants differed from those observed for LOF variants. Homozygous DEL load increased with F_ROH_, consistent with greater exposure of mildly deleterious derived alleles as a larger fraction of the genome becomes autozygous. DEL variants were also relatively enriched within ROHs in both mainland and island populations (Figure 4C, D). Because many ROH regions are shared among individuals and appear to reflect older demographic processes rather than only recent consanguinity, this enrichment may reflect genomic regions in which local reductions in effective population size have both increased homozygosity and reduced the efficacy of purifying selection (Chen et al., 2025; Khan et al., 2021; Xie et al., 2022). However, variation in recombination rate, background selection, genetic draft, and local genomic composition could also contribute to this pattern.

Taken together, these results are compatible with stronger removal of some highly deleterious variants, while mildly deleterious variation persisted or accumulated in genomic regions characterised by elevated homozygosity and reduced effective population size. Because the same demographic and genomic processes that generate persistent ROHs can also alter deleterious-allele frequencies, these patterns should be regarded as suggestive rather than definitive evidence of purging. The combination of reduced diversity and elevated genetic load may nevertheless increase the vulnerability of island populations to future environmental and demographic perturbations, particularly if further inbreeding exposes remaining recessive deleterious variation (Frankham, 1998; Tanaka, 1997; Wright et al., 2008).

### Genomic basis of inbreeding-associated variation in bite force

The combined effects of genetic load, extensive autozygosity, and limited removal of weakly deleterious mutations may leave insular *P. siculus* populations vulnerable to inbreeding depression (Hedrick & Garcia-Dorado, 2016; Keller & Waller, 2002). We therefore examined whether this genomic burden was reflected in whole-organism performance.

Across island populations, individuals with greater genomic autozygosity (F_ROH_) had significantly lower bite force, a pattern consistent with inbreeding depression in this performance trait (Figure 5). Importantly, this negative relationship remained significant when F_ROH_ was centred within populations, confirming that the association was present within island populations and not solely driven by differences among them. Consistent with this result the F_ROH_-bite force relationship was negative within each of the three island populations, although the population-specific slopes were individually uncertain given the moderate sample sizes (Figure S4). Together, these complementary analyses provide consistent evidence that greater genomic inbreeding is associated with reduced bite performance within insular *P. siculus* populations.

A relationship between genomic homozygosity and bite performance was further supported by testing associations between ROHs at individual genomic regions and bite force. Eleven of the thirteen significant windows in island populations were associated with reduced rather than increased bite force, and inspection of the population-specific distributions showed that significant associations reflected segregation of ROHs among individuals within populations rather than by differences among island populations (Figures 6, S5). Most were driven by variation within Sušac, with additional windows segregating within Pod Kopište or across multiple populations. Thus, the genome-wide association between autozygosity and reduced bite force was accompanied by associations at specific genomic regions generated by within-population variation. Among the eleven genes overlapping negatively associated windows, SCN8A and IGDCC3 are involved in cranial skeletomuscular development and function (De Repentigny et al., 2001; Franchini et al., 2019; Porter et al., 1996; W. Yang et al., 2001), making them candidates for contributing to variation in bite performance, although these functional interpretations remain speculative.

Bite force influences competitive ability, prey exploitation, and reproductive success in *Podarcis* (Herrel et al., 2001; Huyghe et al., 2005; Lappin & Husak, 2005; Verwaijen et al., 2002), suggesting that reduced performance in this key functional trait could have fitness consequences. However, the phenotypic consequences of genomic erosion are likely to depend strongly on ecological context and on the extent to which plasticity or adaptation can maintain performance. Pod Mrčaru provides a striking example: despite reduced genetic diversity, lizards from this partly herbivorous population exhibit relatively high size-corrected bite force (Herrel et al., 2008; Taverne et al., 2023). Previous studies suggest that this ecological shift has been facilitated by both phenotypic plasticity and adaptive evolution (Sabolić et al., 2024; Sherpa, Paris, et al., 2023), raising the possibility that ecological and phenotypic flexibility can partly offset the functional consequences of genomic erosion. More generally, this suggests that genomic load need not translate uniformly into reduced performance, and that its phenotypic effects may depend on the ecological and evolutionary context in which populations persist.

In contrast to bite force, we detected no evidence of an association between F_ROH_ and sprint speed, sperm quality, immunocompetence, or spatial cognition. This absence of detectable effects may reflect limited statistical power, differences in trait repeatability and measurement error, or differences in genetic architecture among traits. Weak effects are particularly difficult to detect in moderately sized samples and for traits with substantial environmental or phenotypically plastic variation (Ponzi et al., 2018; Kalinowski & Hedrick, 1999; Kardos et al., 2016). Consequently, the lack of significant associations should not be interpreted as evidence that these traits are unaffected by inbreeding, particularly if their underlying effects are weak, heterogeneous, or genetically diffuse.

The strength and detectability of inbreeding effects also depend on the frequencies and effect sizes of the underlying deleterious variants (Kardos et al., 2016). Rare deleterious variants may have biologically relevant effects without producing a detectable relationship between genome-wide autozygosity and performance, potentially explaining the absence of a F_ROH_-bite force association in mainland lizards despite significant associations with particular segregating ROHs (cf. Duntsch et al., 2023). In small island populations, genetic drift can increase the frequencies of deleterious alleles and associated homozygous haplotypes, potentially bringing their phenotypic effects into a frequency range where they become detectable through among-individual variation mapping. However, this detectability is lost again once such regions become fixed, such as the approximately 2% of the genome we found fixed in ROH across island populations (Figure S3). Phenotypic effects arising from such fixed homozygous regions cannot be mapped through within-population association analyses, meaning that some components of the genetic burden associated with insularity may remain undetectable once the underlying regions have become fixed.

The trait-specific differences in detectable inbreeding effects may also reflect variation in the effect-size distribution and past selective removal of deleterious variants. Our results are consistent with partial removal of strongly deleterious variation in island populations, whereas more weakly deleterious variants appear to have persisted. The widespread distribution of ROH-bite force associations (Figure 6), together with evidence for a highly polygenic architecture of bite force in *P. siculus* (Sabolić et al., 2024), is consistent with contributions from many genomic regions whose individual effects may be relatively modest and therefore less efficiently purged (Bersabé & García-Dorado, 2013). More generally, differences among traits in the frequencies and effect sizes of causal deleterious variants, as well as in their history of purging, could influence how readily inbreeding effects are detected. This may help explain why bite force, but not the other traits examined, showed a detectable association with genomic inbreeding.

To conclude, our results demonstrate that the demographic history of insular *P. siculus* populations has left a clear genomic signature of reduced diversity, increased inbreeding, and elevated genetic load, with evidence for purging of strongly but not mildly deleterious variation. This genomic burden was associated with reduced bite force in more inbred island individuals, whereas other functionally important traits showed no detectable relationship with inbreeding, highlighting trait-specific consequences of genomic erosion. Phenotypic and ecological flexibility may buffer these effects in some contexts, suggesting that the consequences of increased genetic load for population performance may depend on both the trait and the environment.

## Materials and Methods

### Study species

The Italian wall lizard*, Podarcis siculus*, is widely distributed across mainland Italy and numerous Mediterranean islands in the Adriatic and Tyrrhenian seas (Gallozzi et al., 2023; Podnar et al., 2005; Senczuk et al., 2017, 2026; Sherpa, Salvi, et al., 2023). Populations differ markedly in demographic history due to both natural colonisation and human-mediated dispersal (Podnar et al., 2005; Sherpa, Salvi, et al., 2023), including recently founded island populations such as Pod Mrčaru (Croatia), which was established by ten individuals introduced in 1971 and subsequently underwent rapid phenotypic and genomic divergence from its source population (Herrel et al., 2008; Nevo et al., 1972; Sabolić et al., 2024; Sherpa, Paris, et al., 2023; Taverne et al., 2023; Vervust et al., 2007; Wehrle et al., 2020).

### Sample collection, sequencing, and genotyping

Tail tips (∼0.5 cm) were collected from 263 adult Italian wall lizards (187 males and 65 females; mean SVL=70.13 mm, range=47.63-97.26 mm) captured by noose or baited pitfall traps in thirteen populations across the central Adriatic region between September 2022 and May 2024 (Figure 1A). Sampling included three populations from mainland Italy (Termoli, TM, N=40; Rodi Garganico, RG, N=16; Vieste, VS, N=38), four from islands in the Lastovo archipelago, Croatia (Sušac, SC, N=43; Kopište, KP, N=13; Pod Kopište, PK, N=46; Pod Mrčaru, PM, N=46), and one from the Croatian mainland (Trogir, TR, N=5). We additionally included tissue samples from two Croatian mainland populations collected in 2004 (Split, SP, N=2; Paklenica, PL, N=3) and three Croatian island populations collected in 2016 (Mala Palagruža, MP, N=3; Bijelac, BJ, N=5; Pijavica, PJ, N=3). Part of the sequencing data was previously published in Van Linden et al (2025). Sampling locations, years, and sample sizes are summarised in Figure 1A and Table S1. Sampling was conducted under permits issued by Italian Ministry of the Environment and Energy Security (permit numbers: 14060/PNM, 74561/PNM) and the Croatian Ministry of Economy and Sustainable Development (permit numbers: 517-10-1-2-23-4, 517-10-1-2-24-2).

Tail tissue was preserved in 100% ethanol and stored frozen until processing. DNA was extracted using the DNeasy Blood & Tissue Kit (QIAGEN). Whole-genome genomic libraries were prepared and sequenced (paired-end 150 bp) on a DNBSEQ platform by BGI (Shenzhen, China). Read quality was assessed with FASTQC v0.11.9 and summarised using MultiQC v1.12 (Ewels et al., 2016).

Reads were aligned to the *P. siculus* reference genome (rPodSic1.hap1.1; assembly accession GCA_964188175.1) using BWA-MEM v0.7.17 (Li & Durbin, 2009). Resulting BAM files were sorted, indexed, and quality-checked with SAMtools v1.14 (Li et al., 2009). Variants were called using BCFtools v1.14 (Danecek et al., 2021) and filtered to remove variant sites where >10% of reads had mapping quality 0 or where overall mapping quality was <50. Additional filters excluded loci with sequencing depth deviating by more than 2.8 standard deviations from the mean across samples or showing reference/alternative allele read depth bias (Phred >20). Only biallelic SNPs with <20% missing genotypes were retained for downstream analyses.

### Population genomic analyses

Population genetic structure was investigated using PLINK v2.0 (Chang et al., 2015). SNPs in linkage disequilibrium were pruned using a sliding-window approach (50 SNP window, 10 SNP step) removing SNP pairs with r² >0.2 SNPs and with minor allele frequency <0.01. Principal component analyses (PCA) were performed on the pruned dataset for all samples and for island populations separately. We further inferred genetic clustering using ADMIXTURE v1.3.0 (Alexander et al., 2009) for K=1-13, using the PLINK binary format as input. Each analysis included 50 bootstrap replicates and 5-fold cross-validation, with the optimal K determined by the lowest cross-validation error.

Phylogenetic relationships among *P. siculus* populations were inferred using a genomic window-based approach. Non-overlapping 100 kb windows were extracted from the unpruned multi-sample VCF using BCFtools v1.14 (Danecek et al., 2021) and converted to PHYLIP format with vcf2phylip v2.0 (Ortiz, 2019). Maximum likelihood trees were reconstructed for each window with IQ-TREE v2.3.6 (Minh et al., 2020) under a GTR model with 1000 bootstrap replicates, using *Podarcis muralis* (accession SRR8468530) as the outgroup. Window trees were combined into a consensus species tree using ASTRAL-III (Rabiee et al., 2019).

Gene flow among populations was assessed using Patterson’s D-statistic (ABBA-BABA test) implemented in Dsuite (Dtrios, outgroup *P. muralis*; Malinsky et al., 2021). Genome-wide mean F_ST_ between mainland and island populations was calculated in VCFtools (Danecek et al., 2011) using a sliding-window approach (50 kb windows, 10 kb step).

### Demographic history inference

Historical effective population size *N_e_* trajectories were inferred using the Bayesian coalescent framework implemented in PHLASH (Terhorst, 2025). VCF files were first ancestral-state polarised using *P. muralis* as an outgroup (polarizeVCFbyOutgroup.py; Ullrich, 2025). Analyses were performed separately for each island population and for the Italian (IT) and Croatian (HR) mainland populations. The mutation rate was set to 3.96e^-09^ per base per generation, following estimates from *Zootoca vivipara* (Yurchenko et al., 2020). Posterior *N_e_* trajectories were summarised on a log-spaced time grid, assuming a generation time of two years (W. Yang et al., 2021).

### Genomic diversity and inbreeding

Genome-wide heterozygosity, runs of homozygosity (ROHs), and inbreeding coefficients were estimated for each individual. Heterozygosity was inferred from BAM files using ANGSD v0.941 (Korneliussen et al., 2014). Genotype likelihoods were calculated using the SAMtools model with minimum mapping and base quality thresholds of 30 and 20, respectively. Folded site frequency spectra were estimated with realSFS (Nielsen et al., 2012) in non-overlapping 100 kb windows, and heterozygosity was calculated as the proportion of heterozygous sites inferred from genotype likelihoods and averaged across the genome.

ROHs were inferred with BCFtools/RoH (Narasimhan et al., 2016) which uses a hidden Markov model. Transition probabilities between autozygous and non-autozygous states were first estimated using the Viterbi algorithm and subsequently applied for ROH detection (--hw-to-az 0.000624, --az-to-hw 0.00199). Individual inbreeding coefficients based on ROHs (F_ROH_) were calculated as the total length of ROHs divided by the autosomal genome length (McQuillan et al., 2008). ROH length distributions were used to infer the timing of inbreeding events, with longer ROHs indicating more recent inbreeding (Kirin et al., 2010). ROHs were grouped into length classes (10-50 kb, 50-100 kb, 100-500 kb, 500-1000 kb, >1000 kb), and the number of generations since inbreeding was approximated following Thompson (2013): 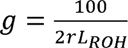, using a recombination rate of ∼1.6 cM/Mb as estimated for *Z. vivipara* (Yurchenko et al., 2020). Differences in genome-wide heterozygosity and F_ROH_, as well as their relationship, and ROH counts per bin, between mainland and island populations were tested using (generalised) linear mixed models (GLMER, LMER) with population as a random effect with the R package *lmerTest* (Kuznetsova et al., 2017) in R v4.4.0 (R Core Team, 2024).

Individual inbreeding coefficients, reflecting deviations from Hardy-Weinberg expectations in observed heterozygosity (F), were estimated using the --het function in VCFtools (Danecek et al., 2011) and averaged per population to obtain population-level estimates of contemporary inbreeding.

### Genetic load

Genetic load of individuals and populations was estimated using an annotation-based approach. As no annotation exists for *P. siculus*, gene models were transferred from *P. muralis* using Liftoff v1.6.3 (-copies, -sc 0.95; Shumate & Salzberg, 2021). A SnpEff database was built from these annotations, and variants in the polarised VCF (ancestral alleles inferred using *P. muralis* as outgroup) were functionally annotated with SnpEff (default parameters; Cingolani et al., 2012).

Variants were classified as synonymous (SYN), missense (DEL), or loss-of-function (LOF; including stop-gain, stop-loss, and splice-site variants), corresponding to SnpEff LOW, MODERATE, and HIGH impact categories (Cingolani et al., 2012). For each individual, the total number of deleterious alleles (weighted additively: heterozygotes=1, homozygotes=2), and the number of heterozygous and homozygous derived genotypes were counted per category. Genetic load was estimated as DEL/SYN and LOF/SYN ratios, calculated genome-wide for heterozygous and homozygous genotypes to reflect masked and realised load, respectively, and for total load separately within ROH and non-ROH regions (Bertorelle et al., 2022).

Allele frequency distributions were assessed using the unfolded site frequency spectrum (SFS). Variants were binned by derived allele frequency, and within each bin the proportions of SYN, DEL, and LOF variants and nonsynonymous to synonymous mutation ratios (NS/S) were calculated.

Differences in genetic load (natural log-transformed) between mainland and island populations, quantified as DEL/SYN and LOF/SYN ratios, and its association with individual genomic inbreeding (F_ROH_) were tested using linear mixed models (LMER) with population as a random effect, using R package *lmerTest* (Kuznetsova et al., 2017) in R v4.4.0 (R Core Team, 2024). For the association with F_ROH_, the models additionally included population origin, the F_ROH_ × origin interaction, and the log-transformed number of homozygous synonymous variants as a covariate.

### Performance measurements and inbreeding depression

We quantified inbreeding depression in mainland and island populations, focusing on performance traits known to be closely linked to fitness in *Podarcis* lizards and potentially sensitive to the deleterious effects of inbreeding, including bite force, sprint speed, sperm quality, immunocompetence, and spatial learning ability (De Meester et al., 2022; Gavriilidi & Van Linden, 2024; Lappin & Husak, 2005; Losdat et al., 2014; Zamora-Camacho et al., 2015). Performance data were collected during the 2024 reproductive season for a subset of 134 adult males from two mainland (VS, N=26; TM, N=28) and three island populations (SC, N=26; PK, N=27; PM, N=27). Only males were tested due to strong sexual dimorphism and to avoid gravid females during the breeding season, while maximizing sample size. In short, bite force was measured using an isometric force transducer (Herrel et al., 1999) and maximal sprint speed was recorded on a 2 m racetrack equipped with photocells (Vanhooydonck et al., 2001). Sperm samples were obtained by abdominal massage and analysed for sperm velocity, count, and morphology (Kahrl et al., 2021). Immunocompetence was assessed using a phytohaemagglutinin (PHA) swelling assay (Mugabo et al., 2010), and spatial cognition was quantified using a spatial learning task (Gavriilidi et al., 2026). Parasite loads were also quantified to account for variation in immune condition (Garrido & Pérez-Mellado, 2014). Detailed experimental procedures are provided in the Supplementary Methods. All procedures were approved by the University of Antwerp’s ethical committee (2023-04 (B)) and by the Italian Ministry of the Environment and Energy Security (permit numbers: 14060/PNM, 74561/PNM) and the Croatian Ministry of Economy and Sustainable Development (permit numbers: 517-10-1-2-23-4, 517-10-1-2-24-2).

To test for inbreeding depression, we modelled the relationship between genomic inbreeding (F_ROH_) and performance traits using Bayesian generalised linear mixed models implemented in *MCMCglmm* (Hadfield, 2024) in R v4.4.0 (R Core Team, 2024). Population origin (mainland vs. island), F_ROH_, and their interaction were included as fixed effects, and population and a centred genomic relatedness matrix (GRM) (estimated with GEMMA; option -gk 1, Zhou and Stephens 2012) were included as random effects to account for population structure (Lavanchy, Weir, et al., 2024). Body size (natural log-transformed SVL) and additional trait-specific covariates were included where appropriate (see Supplementary Results for full model specifications per trait, including covariates and response distributions, Table S7-13). For bite force, we additionally tested whether the F_ROH_ association persisted within island populations by centring F_ROH_ around each population mean, and tested for differences in F_ROH_ slopes among island populations by including F_ROH_ × population interactions. Models were fitted with non-informative inverse-Wishart priors and run sufficiently long to obtain an effective sample size greater than ∼9000 for the main model parameters (2.000.000 MCMC iterations; burn-in 200.000; thinning 200). Convergence was assessed using trace plots, autocorrelation, and effective sample sizes. Effects were evaluated using posterior means (β), 95% credible intervals, and pMCMC values.

### ROH-GWAS: genetic architecture of inbreeding depression

We performed a genome-wide association study of window-based runs of homozygosity (ROH-GWAS) with performance traits that showed significant inbreeding depression in the analysis described above, following frameworks described by Stoffel et al. (2021), Duntsch et al. (2023), and Hewett et al. (2026). The genome was partitioned into non-overlapping windows of 1500 SNPs (∼16.5 kb, SNP density of 1 SNP/11 bp). For each individual, a window was considered in ROH if ≥50% of SNPs overlapped a ROH detected with BCFtools/RoH (ROH_status_=1). These windowed ROH genome landscapes were summarised for mainland and island groups, and similarity in landscape among populations and between groups was quantified using pairwise Pearson correlations. For the ROH-GWAS, we tested for each window whether ROH presence was associated with the standardized phenotype using linear models including ROH_status_, genome-wide inbreeding (F_ROH_ excluding the focal chromosome, to control for overall level of inbreeding), and principal components from the GRM (PCs, to control for population structure). Analyses were performed separately for mainland and island populations. Windows were considered non-informative and excluded from analysis when fewer than 20 individuals had complete data or when fewer than three individuals were present in either the ROH or non-ROH group. Multiple testing correction was applied using the effective number of independent tests (N_eff_) estimated with the simpleM approach (Gao et al., 2008), to account for strong correlation among windows because of long ROH tracts. Significant windows were inspected to identify candidate genes by intersecting ROH windows with gene coordinates derived from the Liftoff annotation (BEDTools intersect, Quinlan & Hall, 2010).

## Acknowledgements

The authors thank Jan Scholliers, Jorrit Mertens, Simon Baeckens, and the staff of Nature Park Lastovo Islands for help with the fieldwork. Financial support was given by Research Foundation Flanders (FWO) grant 1102623N to LVL and by University of Antwerp to LVL, RVD, HS. Sample and data collection was funded by FWO grant V430624N to LVL, Croatian Science Foundation grant HRZZ-IP-06-2016-9177 to AŠ, and National Geographic Committee for Research and Exploration to AH.

## Data Accessibility Statement

All raw sequencing data are available on NCBI and SRA under BioProject ID PRJNA1225591 (released upon publication). Scripts for sequence alignment, variant calling, filtering, population genomics, genetic load, and inbreeding depression analyses are available at https://github.com/lisavanlinden/P.siculus_genetic_load.git.

## Author Contributions

RVD, HS, and LVL originally formulated the idea. AH and AŠ provided samples from 2016 and AŠ sampling permits for 2022-2024. LVL, ZM, IG, and RVD conducted the fieldwork in 2022-2024, including sampling and experimental procedures. LVL, ZM, and IG collected and extracted the performance data and conducted the (preliminary) analyses. LVL analysed the data with input from HS, RVD, and IG. CJF conducted the demographic history analyses. LVL wrote the first draft, with substantial input from HS. All authors contributed to the final manuscript.

## References

Alexander, D. H., Novembre, J., & Lange, K. (2009). Fast model-based estimation of ancestry in unrelated individuals. Genome Research, 19(9), 1655. 10.1101/gr.094052.109

Allendorf, F. W., Funk, W. C., Aitken, S. N., Byrne, M., & Luikart, G. (2022). Conservation and the genomics of populations (3rd ed.). Oxford University Press.

Arnold, S. J. (1983). Morphology, performance and fitness. American Zoologist, 23(2), 347–361.

Bersabé, D., & García-Dorado, A. (2013). On the genetic parameter determining the efficiency of purging: An estimate for *Drosophila* egg-to-pupae viability. Journal of Evolutionary Biology, 26(2), 375–385. 10.1111/jeb.12054

Bertorelle, G., Raffini, F., Bosse, M., Bortoluzzi, C., Iannucci, A., Trucchi, E., Morales, H. E., & van Oosterhout, C. (2022). Genetic load: Genomic estimates and applications in non-model animals. Nature Reviews Genetics, 23(8), 492–503. 10.1038/s41576-022-00448-x

Biaggini, M., & Corti, C. (2019). Notes on egg and hatchling size in *Podarcis siculus* (Squamata: Lacertidae) from central Italy. Phyllomedusa: Journal of Herpetology, 18(1), Article 1. 10.11606/issn.2316-9079.v18i1p127-129

Böhme, M. U., Fritz, U., Kotenko, T., Džukić, G., Ljubisavljević, K., Tzankov, N., & Berendonk, T. U. (2007). Phylogeography and cryptic variation within the *Lacerta viridis* complex (Lacertidae, Reptilia). Zoologica Scripta, 36(2), 119–131. 10.1111/j.1463-6409.2006.00262.x

Bortoluzzi, C., Bosse, M., Derks, M. F. L., Crooijmans, R. P. M. A., Groenen, M. A. M., & Megens, H.-J. (2020). The type of bottleneck matters: Insights into the deleterious variation landscape of small managed populations. Evolutionary Applications, 13(2), 330–341. 10.1111/eva.12872

Brewer, B. A., Lacy, R. C., Foster, M. L., & Alaks, G. (1990). Inbreeding depression in insular and central populations of *Peromyscus* mice. Journal of Heredity, 81(4), 257–266. 10.1093/oxfordjournals.jhered.a110988

Canestrelli, D., Salvi, D., Maura, M., Bologna, M. A., & Nascetti, G. (2012). One species, three Pleistocene evolutionary histories: Phylogeography of the Italian crested newt, *Triturus carnifex*. PLOS ONE, 7(7), e41754. 10.1371/journal.pone.0041754

Carretero, M. A. (2006). Reproductive cycles in Mediterranean lacertids: Plasticity and constraints. In Mainland and insular lacertid lizards: A Mediterranean perspective. Edited by C. Corti, P. Lo Cascio, M. Biaggini (pp. 33–54). Firenze University Press.

Cavill, E. L., Morales, H. E., Sun, X., Westbury, M. V., van Oosterhout, C., Accouche, W., Zora, A., Schulze, M. J., Shah, N., Adam, P.-A., Brooke, M. de L., Sweet, P., Gopalakrishnan, S., & Gilbert, M. T. P. (2024). When birds of a feather flock together: Severe genomic erosion and the implications for genetic rescue in an endangered island passerine. Evolutionary Applications, 17(7), e13739. 10.1111/eva.13739

Chang, C. C., Chow, C. C., Tellier, L. C., Vattikuti, S., Purcell, S. M., & Lee, J. J. (2015). Second-generation PLINK: Rising to the challenge of larger and richer datasets. GigaScience, 4(1), s13742–015-0047–0048. 10.1186/s13742-015-0047-8

Charlesworth, D., & Willis, J. H. (2009). The genetics of inbreeding depression. Nature Reviews Genetics, 10(11), Article 11. 10.1038/nrg2664

Chen, Y., Feng, X., Reid, K., Zhang, C., Löytynoja, A., & Merilä, J. (2025). Dynamics of deleterious mutations and purifying selection in small population isolates. Molecular Biology and Evolution, 42(7). 10.1093/molbev/msaf110

Cingolani, P., Platts, A., Wang, L. L., Coon, M., Nguyen, T., Wang, L., Land, S. J., Lu, X., & Ruden, D. M. (2012). A program for annotating and predicting the effects of single nucleotide polymorphisms, SnpEff. Fly, 6(2), 80–92. 10.4161/fly.19695

Crow, J. F. (James F. (with Internet Archive). (1970). An introduction to population genetics theory. New York, Harper & Row. http://archive.org/details/introductiontopo0000crow

Danecek, P., Auton, A., Abecasis, G., Albers, C. A., Banks, E., DePristo, M. A., Handsaker, R. E., Lunter, G., Marth, G. T., Sherry, S. T., McVean, G., Durbin, R., & 1000 Genomes Project Analysis Group. (2011). The variant call format and VCFtools. Bioinformatics, 27(15), 2156–2158. 10.1093/bioinformatics/btr330

Danecek, P., Bonfield, J. K., Liddle, J., Marshall, J., Ohan, V., Pollard, M. O., Whitwham, A., Keane, T., McCarthy, S. A., Davies, R. M., & Li, H. (2021). Twelve years of SAMtools and BCFtools. GigaScience, 10(2), giab008. 10.1093/gigascience/giab008

De Meester, G., Van Linden, L., Torfs, J., Pafilis, P., Šunje, E., Steenssens, D., Zulčić, T., Sassalos, A., & Van Damme, R. (2022). Learning with lacertids: Studying the link between ecology and cognition within a comparative framework. Evolution, 76(11), 2531–2552. 10.1111/evo.14618

De Repentigny, Y., Côté, P. D., Pool, M., Bernier, G., Girard, S., Vidal, S. M., & Kothary, R. (2001). Pathological and genetic analysis of the degenerating muscle (dmu) mouse: A new allele of Scn8a. Human Molecular Genetics, 10(17), 1819–1827. 10.1093/hmg/10.17.1819

Duntsch, L., Whibley, A., de Villemereuil, P., Brekke, P., Bailey, S., Ewen, J. G., & Santure, A. W. (2023). Genomic signatures of inbreeding depression for a threatened Aotearoa New Zealand passerine. Molecular Ecology, 32(8), 1893–1907. 10.1111/mec.16855

Dussex, N., Morales, H. E., Grossen, C., Dalén, L., & Oosterhout, C. van. (2023). Purging and accumulation of genetic load in conservation. Trends in Ecology & Evolution, 0(0). 10.1016/j.tree.2023.05.008

Dussex, N., van der Valk, T., Morales, H. E., Wheat, C. W., Díez-del-Molino, D., von Seth, J., Foster, Y., Kutschera, V. E., Guschanski, K., Rhie, A., Phillippy, A. M., Korlach, J., Howe, K., Chow, W., Pelan, S., Mendes Damas, J. D., Lewin, H. A., Hastie, A. R., Formenti, G., … Dalén, L. (2021). Population genomics of the critically endangered kākāpō. Cell Genomics, 1(1), 100002. 10.1016/j.xgen.2021.100002

Eldridge, M. D. B., King, J. M., Loupis, A. K., Spencer, P. B. S., Taylor, A. C., Pope, L. C., & Hall, G. P. (1999). Unprecedented Low Levels of Genetic Variation and Inbreeding Depression in an Island Population of the Black-Footed Rock-Wallaby. Conservation Biology, 13(3), 531–541. 10.1046/j.1523-1739.1999.98115.x

Ewels, P., Magnusson, M., Lundin, S., & Käller, M. (2016). MultiQC: Summarize analysis results for multiple tools and samples in a single report. Bioinformatics, 32(19), 3047–3048. 10.1093/bioinformatics/btw354

Franchini, P., Xiong, P., Fruciano, C., Schneider, R. F., Woltering, J. M., Hulsey, C. D., & Meyer, A. (2019). MicroRNA gene reulation in extremely young and parallel adaptive radiations of crater lake cichlid fish. Molecular Biology and Evolution, 36(11), 2498–2511. 10.1093/molbev/msz168

Frankham, R. (1998). Inbreeding and extinction: Island populations. Conservation Biology, 12(3), 665–675. 10.1111/j.1523-1739.1998.96456.x

Gabrielli, M., Benazzo, A., Biello, R., Iannucci, A., Salvi, D., Ficetola, G. F., Ciofi, C., Trucchi, E., & Bertorelle, G. (2026). The relationship between genomic variation and genetic load: Insights from small island populations. Heredity, 1–10. 10.1038/s41437-026-00835-8

Gabrielli, M., Morales, H. E., Noguerales, V., Wei, W., Cotoras, D. D., Dussex, N., Pereira, R. J., Emerson, B. C., & Cerca, J. (2026). The paradox of persistence: Island species as models for conservation. Trends in Ecology & Evolution. 10.1016/j.tree.2026.08.006

Gallozzi, F., Corti, C., Castiglia, R., Avramo, V., Senczuk, G., Mattioni, C., & Colangelo, P. (2023). The intriguing biogeographic pattern of the Italian wall lizard *Podarcis siculus* (Squamata: Lacertidae) in the Tuscan archipelago reveals the existence of a new ancient insular clade. Animals, 13(3), Article 3. 10.3390/ani13030386

Gao, X., Starmer, J., & Martin, E. R. (2008). A multiple testing correction method for genetic association studies using correlated single nucleotide polymorphisms. Genetic Epidemiology, 32(4), 361–369. 10.1002/gepi.20310

García-Verdugo, C., Sajeva, M., La Mantia, T., Harrouni, C., Msanda, F., & Caujapé-Castells, J. (2015). Do island plant populations really have lower genetic variation than mainland populations? Effects of selection and distribution range on genetic diversity estimates. Molecular Ecology, 24(4), 726–741. 10.1111/mec.13060

Gargiulo, R., Budde, K. B., & Heuertz, M. (2024). Mind the lag: Understanding genetic extinction debt for conservation. Trends in Ecology & Evolution, 0(0). 10.1016/j.tree.2024.10.008

Garrido, M., & Pérez-Mellado, V. (2014). Immune responses vary with parasite burden in an insular lizard. Evolutionary Ecology Research, 16, 593–603.

Gavriilidi, I., Baeckens, S., Pafilis, P., & Van Damme, R. (2026). Creatures of habit: Similar but more repeatable behaviours in insular compared with mainland lizards. Animal Behaviour, 233, 123493. 10.1016/j.anbehav.2026.123493

Gavriilidi, I., & Van Linden, L. (2024). Inbreeding and cognition in wild populations: A relationship that remains unnoticed. Oikos, 2024(9), e10674. 10.1111/oik.10674

Giovannotti, M., Nisi-Cerioni, P., & Caputo, V. (2010). Mitochondrial DNA sequence analysis reveals multiple Pleistocene glacial refugia for *Podarcis muralis* (Laurenti, 1768) in the Italian Peninsula. Italian Journal of Zoology, 77(3), 277–288. 10.1080/11250000903143885

Hadfield, J. (2024). MCMCglmm: MCMC Generalised Linear Mixed Models (Version 2.36) [Computer software]. https://cran.r-project.org/web/packages/MCMCglmm/index.html

Hasselgren, M., Dussex, N., von Seth, J., Angerbjörn, A., Dalén, L., & Norén, K. (2024). Strongly deleterious mutations influence reproductive output and longevity in an endangered population. Nature Communications, 15(1), 8378. 10.1038/s41467-024-52741-4

Hedrick, P. W., & Garcia-Dorado, A. (2016). Understanding inbreeding depression, purging, and genetic rescue. Trends in Ecology & Evolution, 31(12), 940–952. 10.1016/j.tree.2016.09.005

Hedrick, P. W., & Kalinowski, S. T. (2000). Inbreeding depression in conservation biology. Annual Review of Ecology and Systematics, 31(1), 139–162. 10.1146/annurev.ecolsys.31.1.139

Herrel, A., Damme, R. V., Vanhooydonck, B., & Vree, F. D. (2001). The implications of bite performance for diet in two species of lacertid lizards. Canadian Journal of Zoology, 79(4), 662–670. 10.1139/z01-031

Herrel, A., Huyghe, K., Vanhooydonck, B., Backeljau, T., Breugelmans, K., Grbac, I., Van Damme, R., & Irschick, D. J. (2008). Rapid large-scale evolutionary divergence in morphology and performance associated with exploitation of a different dietary resource. Proceedings of the National Academy of Sciences, 105(12), 4792–4795. 10.1073/pnas.0711998105

Herrel, A., Spithoven, L., Van Damme, R., & DE Vree, F. (1999). Sexual dimorphism of head size in *Gallotia galloti*: Testing the niche divergence hypothesis by functional analyses. Functional Ecology, 13(3), 289–297. 10.1046/j.1365-2435.1999.00305.x

Hewett, A., Lavanchy, E., Topaloudis, A., Cumer, T., Ducrest, A.-L., Simon, C., Almasi, B., Roulin, A., & Goudet, J. (2026). Inbreeding depression in morphological and growth traits of wild barn owls (*Tyto alba*). Proceedings of the Royal Society B: Biological Sciences, 293(2070), 20252708. 10.1098/rspb.2025.2708

Hurston, H., Voith, L., Bonanno, J., Foufopoulos, J., Pafilis, P., Valakos, E., & Anthony, N. (2009). Effects of fragmentation on genetic diversity in island populations of the Aegean wall lizard *Podarcis erhardii* (Lacertidae, Reptilia). Molecular Phylogenetics and Evolution, 52(2), 395–405. 10.1016/j.ympev.2009.03.028

Huyghe, K., Vanhooydonck, B., Scheers, H., Molina-Borja, M., & Van Damme, R. (2005). Morphology, performance and fighting capacity in male lizards, *Gallotia galloti*. Functional Ecology, 19(5), 800–807.

James, J. E., Lanfear, R., & Eyre-Walker, A. (2016). Molecular evolutionary consequences of island colonization. Genome Biology and Evolution, 8(6), 1876–1888. 10.1093/gbe/evw120

Ji, X., & Braña, F. (2000). Among clutch variation in reproductive output and egg size in the wall lizard (*Podarcis muralis*) from a lowland population of northern Spain. Journal of Herpetology, 34(1), 54–60. 10.2307/1565238

Kahrl, A. F., Kustra, M. C., Reedy, A. M., Bhave, R. S., Seears, H. A., Warner, D. A., & Cox, R. M. (2021). Selection on Sperm Count, but Not on Sperm Morphology or Velocity, in a Wild Population of Anolis Lizards. Cells, 10(9), Article 9. 10.3390/cells10092369

Kalinowski, S. T., & Hedrick, P. W. (1999). Detecting inbreeding depression is difficult in captive endangered species. Animal Conservation, 2(2), 131–136. 10.1111/j.1469-1795.1999.tb00058.x

Kardos, M., Armstrong, E. E., Fitzpatrick, S. W., Hauser, S., Hedrick, P. W., Miller, J. M., Tallmon, D. A., & Funk, W. C. (2021). The crucial role of genome-wide genetic variation in conservation. Proceedings of the National Academy of Sciences, 118(48), e2104642118. 10.1073/pnas.2104642118

Kardos, M., Luikart, G., & Allendorf, F. W. (2015). Measuring individual inbreeding in the age of genomics: Marker-based measures are better than pedigrees. Heredity, 115(1), 63–72. 10.1038/hdy.2015.17

Kardos, M., Taylor, H. R., Ellegren, H., Luikart, G., & Allendorf, F. W. (2016). Genomics advances the study of inbreeding depression in the wild. Evolutionary Applications, 9(10), 1205–1218. 10.1111/eva.12414

Keller, L. F., & Waller, D. M. (2002). Inbreeding effects in wild populations. Trends in Ecology & Evolution, 17(5), 230–241. 10.1016/S0169-5347(02)02489-8

Khan, A., Patel, K., Shukla, H., Viswanathan, A., van der Valk, T., Borthakur, U., Nigam, P., Zachariah, A., Jhala, Y. V., Kardos, M., & Ramakrishnan, U. (2021). Genomic evidence for inbreeding depression and purging of deleterious genetic variation in Indian tigers. Proceedings of the National Academy of Sciences, 118(49), e2023018118. 10.1073/pnas.2023018118

Kirin, M., McQuillan, R., Franklin, C. S., Campbell, H., McKeigue, P. M., & Wilson, J. F. (2010). Genomic runs of homozygosity record population history and consanguinity. PLOS ONE, 5(11), e13996. 10.1371/journal.pone.0013996

Korneliussen, T. S., Albrechtsen, A., & Nielsen, R. (2014). ANGSD: Analysis of Next Generation Sequencing Data. BMC Bioinformatics, 15(1), 356. 10.1186/s12859-014-0356-4

Kuznetsova, A., Brockhoff, P. B., & Christensen, R. H. B. (2017). lmerTest Package: Tests in Linear Mixed Effects Models. Journal of Statistical Software, 82, 1–26. 10.18637/jss.v082.i13

Lan, T., Li, H., Liu, B., Shi, M., Tian, Y., Sahu, S. K., Cui, L., Dussex, N., Liu, D., Ma, Y., Kong, W., Liu, S., Fan, J., Zhao, Y., Fu, Y., Li, Q., Lin, C., Dalén, L., Liu, H., … Xu, Y. (2025). Revealing extensive inbreeding and less efficient purging of deleterious mutations in wild Amur tigers in China. Journal of Genetics and Genomics, 52(5), 641–649. 10.1016/j.jgg.2024.12.004

Lappin, A. K., & Husak, J. F. (2005). Weapon performance, not size, determines mating success and potential reproductive output in the collared lizard (*Crotaphytus collaris*). The American Naturalist, 166(3), 426–436. 10.1086/432564

Lavanchy, E., Cumer, T., Topaloudis, A., Ducrest, A.-L., Simon, C., Roulin, A., & Goudet, J. (2024). Too big to purge: Persistence of deleterious mutations in island populations of the European barn owl (*Tyto alba*). Heredity, 133(6), 437–449. 10.1038/s41437-024-00728-8

Lavanchy, E., Weir, B. S., & Goudet, J. (2024). Detecting inbreeding depression in structured populations. Proceedings of the National Academy of Sciences, 121(19), e2315780121. 10.1073/pnas.2315780121

Li, H., & Durbin, R. (2009). Fast and accurate short read alignment with Burrows– Wheeler transform. Bioinformatics, 25(14), 1754–1760. 10.1093/bioinformatics/btp324

Li, H., Handsaker, B., Wysoker, A., Fennell, T., Ruan, J., Homer, N., Marth, G., Abecasis, G., Durbin, R., & 1000 Genome Project Data Processing Subgroup. (2009). The Sequence Alignment/Map format and SAMtools. Bioinformatics, 25(16), 2078–2079. 10.1093/bioinformatics/btp352

Losdat, S., Chang, S.-M., & Reid, J. M. (2014). Inbreeding depression in male gametic performance. Journal of Evolutionary Biology, 27(6), 992–1011. 10.1111/jeb.12403

Lozier, J. D., Strange, J. P., & Heraghty, S. D. (2023). Whole genome demographic models indicate divergent effective population size histories shape contemporary genetic diversity gradients in a montane bumble bee. Ecology and Evolution, 13(2), e9778. 10.1002/ece3.9778

Machado, A. P., Topaloudis, A., Cumer, T., Lavanchy, E., Bontzorlos, V., Ceccherelli, R., Charter, M., Kassinis, N., Lymberakis, P., Manzia, F., Ducrest, A.-L., Dupasquier, M., Guex, N., Roulin, A., & Goudet, J. (2022). Genomic consequences of colonisation, migration and genetic drift in barn owl insular populations of the eastern Mediterranean. Molecular Ecology, 31(5), 1375– 1388. 10.1111/mec.16324

Malinsky, M., Matschiner, M., & Svardal, H. (2021). Dsuite—Fast D-statistics and related admixture evidence from VCF files. Molecular Ecology Resources, 21(2), 584–595. 10.1111/1755-0998.13265

Martin, C. A., Sheppard, E. C., Illera, J. C., Suh, A., Nadachowska-Brzyska, K., Spurgin, L. G., & Richardson, D. S. (2023). Runs of homozygosity reveal past bottlenecks and contemporary inbreeding across diverging populations of an island-colonizing bird. Molecular Ecology, 32(8), 1972–1989. 10.1111/mec.16865

McQuillan, R., Leutenegger, A.-L., Abdel-Rahman, R., Franklin, C. S., Pericic, M., Barac-Lauc, L., Smolej-Narancic, N., Janicijevic, B., Polasek, O., Tenesa, A., MacLeod, A. K., Farrington, S. M., Rudan, P., Hayward, C., Vitart, V., Rudan, I., Wild, S. H., Dunlop, M. G., Wright, A. F., … Wilson, J. F. (2008). Runs of homozygosity in European populations. The American Journal of Human Genetics, 83(3), 359–372. 10.1016/j.ajhg.2008.08.007

Michaelides, S. N., Stroud, J. T., Giery, S. T., Smith, B. E., Glor, R. E., & Kolbe, J. J. (2025). Invasion histories differ markedly among the four *Anolis* lizard species introduced to Bermuda. Journal of Herpetology, 59(2), 78–85. 10.1670/23-043

Minh, B. Q., Schmidt, H. A., Chernomor, O., Schrempf, D., Woodhams, M. D., von Haeseler, A., & Lanfear, R. (2020). IQ-TREE 2: New models and efficient methods for phylogenetic inference in the genomic era. Molecular Biology and Evolution, 37(5), 1530–1534. 10.1093/molbev/msaa015

Mugabo, M., Marquis, O., Perret, S., & Le Galliard, J. F. (2010). Immediate and delayed life history effects caused by food deprivation early in life in a short- lived lizard. Journal of Evolutionary Biology, 23(9), 1886–1898. 10.1111/j.1420-9101.2010.02052.x

Narasimhan, V., Danecek, P., Scally, A., Xue, Y., Tyler-Smith, C., & Durbin, R. (2016). BCFtools/RoH: A hidden Markov model approach for detecting autozygosity from next-generation sequencing data. Bioinformatics, 32(11), 1749–1751. 10.1093/bioinformatics/btw044

Nevo, E., Gorman, G., Soulé, M., Yang, S. Y., Clover, R., & Jovanović, V. (1972). Competitive exclusion between insular Lacerta species (Sauria, Lacertidae): Notes on experimental introductions. Oecologia, 10(2), 183–190. 10.1007/BF00347990

Nielsen, R., Korneliussen, T., Albrechtsen, A., Li, Y., & Wang, J. (2012). SNP calling, genotype calling, and sample allele frequency estimation from new-generation sequencing data. PLOS ONE, 7(7), e37558. 10.1371/journal.pone.0037558

Oppliger, A., Degen, L., Bouteiller-Reuter, C., & John-Alder, H.-B. (2007). Promiscuity and high level of multiple paternity in common wall lizards (*Podarcis muralis*): Data from microsatellite markers. Amphibia-Reptilia, 28(2), 301–303. 10.1163/156853807780202477

Ortiz, E. M. (2019). vcf2phylip v2.0: Convert a VCF matrix into several matrix formats for phylogenetic analysis. [Computer software]. Zenodo. 10.5281/zenodo.2540861

Otalora, K., Riera, J. L., Tavecchia, G., Rotger, A., Igual, J. M., Trotta, J.-R. P., & Baldo, L. (2024). Population genetics and phylogeographic history of the insular lizard *Podarcis lilfordi* (Gunther, 1874) from the Balearic Islands based on genome-wide polymorphic data. Ecology and Evolution, 14(5), e11407. 10.1002/ece3.11407

Plećaš, D., Gotovac Đogaš, V., Polašek, O., & Škunca Herman, J. (2024). Determinants of human asymmetry: Does asymmetrical retinal vasculature predict asymmetry elsewhere in the body? Life, 14(8), 929. 10.3390/life14080929

Podnar, M., Mayer, W., & Tvrtković, N. (2004). Mitochondrial phylogeography of the Dalmatian wall lizard, *Podarcis melisellensis* (Lacertidae). Organisms Diversity & Evolution, 4(4), 307–317. 10.1016/j.ode.2004.04.004

Podnar, M., Mayer, W., & Tvrtković, N. (2005). Phylogeography of the Italian wall lizard, *Podarcis sicula*, as revealed by mitochondrial DNA sequences. Molecular Ecology, 14(2), 575–588. 10.1111/j.1365-294X.2005.02427.x

Ponzi, E., Keller, L. F., Bonnet, T., & Muff, S. (2018). Heritability, selection, and the response to selection in the presence of phenotypic measurement error: Effects, cures, and the role of repeated measurements. Evolution, 72(10), 1992–2004. 10.1111/evo.13573

Porter, J. D., Goldstein, L. A., Kasarskis, E. J., Brueckner, J. K., & Spear, B. T. (1996). The neuronal voltage-gated sodium channel, Scn8a, is essential for postnatal maturation of spinal, but not oculomotor, motor units. Experimental Neurology, 139(2), 328–334. 10.1006/exnr.1996.0107

Quinlan, A. R., & Hall, I. M. (2010). BEDTools: A flexible suite of utilities for comparing genomic features. Bioinformatics, 26(6), 841–842. 10.1093/bioinformatics/btq033

R Core Team. (2024). R: A language and environment for statistical computing. *R Foundation for Statistical Computing*, *Vienna, Austria*. Https://www.R-project.org/. [Computer software]. https://www.R-project.org/.

Rabiee, M., Sayyari, E., & Mirarab, S. (2019). Multi-allele species reconstruction using ASTRAL. Molecular Phylogenetics and Evolution, 130, 286–296. 10.1016/j.ympev.2018.10.033

Robinson, J. A., Brown, C., Kim, B. Y., Lohmueller, K. E., & Wayne, R. K. (2018). Purging of strongly deleterious mutations explains long-term persistence and absence of inbreeding depression in island foxes. Current Biology, 28(21), 3487–3494.e4. 10.1016/j.cub.2018.08.066

Rotger, A., Tenan, S., Igual, J.-M., Bonner, S., & Tavecchia, G. (2023). Life span, growth, senescence and island syndrome: Accounting for imperfect detection and continuous growth. Journal of Animal Ecology, 92(1), 183–194. 10.1111/1365-2656.13842

Runemark, A., Hey, J., Hansson, B., & Svensson, E. I. (2012). Vicariance divergence and gene flow among islet populations of an endemic lizard. Molecular Ecology, 21(1), 117–129. 10.1111/j.1365-294X.2011.05377.x

Sabolić, I. (2021). Genomic patterns of phenotypic adaptation in Italian wall lizard Podarcis siculus (Rafinesque-Schmaltz, 1810) [Doctoral Thesis, University of Zagreb. Faculty of Science. Department of Biology]. https://urn.nsk.hr/urn:nbn:hr:217:896917

Sabolić, I., Mira, Ó., Brandt, D. Y. C., Lisičić, D., Stapley, J., Novosolov, M., Bakarić, R., Cizelj, I., Glogoški, M., Hudina, T., Taverne, M., Allentoft, M. E., Nielsen, R., Herrel, A., & Štambuk, A. (2024). Plastic and genomic change of a newly established lizard population following a founder event. Molecular Ecology, 33(10), e17255. 10.1111/mec.17255

Salvi, D., Harris, D. J., Kaliontzopoulou, A., Carretero, M. A., & Pinho, C. (2013). Persistence across Pleistocene ice ages in Mediterranean and extra-Mediterranean refugia: Phylogeographic insights from the common wall lizard. BMC Evolutionary Biology, 13(1), 147. 10.1186/1471-2148-13-147

Senczuk, G., Colangelo, P., De Simone, E., Aloise, G., & Castiglia, R. (2017). A combination of long term fragmentation and glacial persistence drove the evolutionary history of the Italian wall lizard *Podarcis siculus*. BMC Evolutionary Biology, 17(1), 6. 10.1186/s12862-016-0847-1

Senczuk, G., Ripa, C., Colangelo, P., & Castiglia, R. (2026). Genetic population structure of the Italian wall lizards *Podarcis siculus* (Squamata: Lacertidae): Insight from nuclear DNA markers. Ecology and Evolution, 16(1), e72655. 10.1002/ece3.72655

Sherpa, S., Paris, J. R., Silva-Rocha, I., Di Canio, V., Carretero, M. A., Ficetola, G. F., & Salvi, D. (2023). Genetic depletion does not prevent rapid evolution in island-introduced lizards. Ecology and Evolution, 13(11), e10721. 10.1002/ece3.10721

Sherpa, S., Salvi, D., & Silva-Rocha, I. (2023). Reconstructing the complex colonisation histories of lizards across Mediterranean archipelagos. Journal of Biogeography, 2023, 1–16. 10.1111/jbi.14739

Shumate, A., & Salzberg, S. L. (2021). Liftoff: Accurate mapping of gene annotations. Bioinformatics, 37(12), 1639–1643. 10.1093/bioinformatics/btaa1016

Soulé, M. E. (1987). Viable Populations for Conservation. Cambridge University Press.

Stoffel, M. A., Johnston, S. E., Pilkington, J. G., & Pemberton, J. M. (2021). Genetic architecture and lifetime dynamics of inbreeding depression in a wild mammal. Nature Communications, 12(1), Article 1. 10.1038/s41467-021-23222-9

Tanaka, Y. (1997). Extinction of populations due to inbreeding depression with demographic disturbances. Researches on Population Ecology, 39(1), 57–66. 10.1007/BF02765250

Taverne, M., Watson, P. J., Dutel, H., Boistel, R., Lisicic, D., Tadic, Z., Fabre, A.-C., Fagan, M. J., & Herrel, A. (2023). Form–function relationships underlie rapid dietary changes in a lizard. Proceedings of the Royal Society B: Biological Sciences, 290(2000), 20230582. 10.1098/rspb.2023.0582

Taylor, R. S., Manseau, M., Keobouasone, S., Liu, P., Mastromonaco, G., Solmundson, K., Kelly, A., Larter, N. C., Gamberg, M., Schwantje, H., Thacker, C., Polfus, J., Andrew, L., Hervieux, D., Simmons, D., & Wilson, P. J. (2024). High genetic load without purging in caribou, a diverse species at risk. Current Biology, 34(6), 1234–1246.e7. 10.1016/j.cub.2024.02.002

Terhorst, J. (2025). Accelerated Bayesian inference of population size history from recombining sequence data. Nature Genetics, 57(10), 2570–2577. 10.1038/s41588-025-02323-x

Thompson, E. A. (2013). Identity by Descent: Variation in meiosis, across genomes, and in populations. Genetics, 194(2), 301–326. 10.1534/genetics.112.148825

Tsumura, Y., & Ohba, K. (1993). Genetic structure of geographical marginal populations of *Cryptomeria japonica*. Canadian Journal of Forest Research, 23(5), 859–863. 10.1139/x93-112

Ullrich, K. (2025). polarizeVCFbyOutgroup;py [Computer software]. https://github.com/kullrich/bio-scripts/tree/master/vcf

Van Linden, L., Svardal, H., Štambuk, A., Herrel, A., & Van Damme, R. (2025). Without the locals’ aid: No evidence for a role of admixture in the colonisation success of Italian wall lizards. Oecologia, 207(7), 125. 10.1007/s00442-025-05769-2

Van Noordwijk, A. J., & Scharloo, W. (1981). Inbreeding in an Island Population of the Great Tit. Evolution, 35(4), 674–688. 10.2307/2408240

Vanhooydonck, B., Van Damme, R., Van Dooren, T. J. M., & Bauwens, D. (2001). Proximate causes of intraspecific variation in locomotor performance in the lizard *Gallotia galloti*. Physiological and Biochemical Zoology, 74(6), 937–945. 10.1086/324749

Vervust, B., Grbac, I., & Van Damme, R. (2007). Differences in morphology, performance and behaviour between recently diverged populations of *Podarcis sicula* mirror differences in predation pressure. Oikos, 116(8), 1343– 1352. 10.1111/j.0030-1299.2007.15989.x

Verwaijen, D., Van Damme, R., & Herrel, A. (2002). Relationships between head size, bite force, prey handling efficiency and diet in two sympatric lacertid lizards. Functional Ecology, 16(6), 842–850. 10.1046/j.1365-2435.2002.00696.x

Vlček, J., Espinoza-Ulloa, S., Cowles, S. A., Ortiz-Catedral, L., Coutu, C., Chaves, J. A., Andrés, J., & Štefka, J. (2025). Genomes of Galápagos mockingbirds reveal the impact of island size and past demography on inbreeding and genetic load in contemporary populations. Molecular Ecology, 34(5), e17665. 10.1111/mec.17665

Wehrle, B. A., Herrel, A., Nguyen-Phuc, B.-Q., Maldonado, S., Dang, R. K., Agnihotri, R., Tadić, Z., & German, D. P. (2020). Rapid dietary shift in *Podarcis siculus* resulted in localized changes in gut function. Physiological and Biochemical Zoology, 93(5), 396–415. 10.1086/709848

Whitlock, M., & Davis, B. (2011). Genetic Load. In Wiley, Encyclopedia of Life Sciences (1st ed.). Wiley. 10.1002/9780470015902.a0001787.pub2

Wright, L. I., Tregenza, T., & Hosken, D. J. (2008). Inbreeding, inbreeding depression and extinction. Conservation Genetics, 9(4), 833–843. 10.1007/s10592-007-9405-0

Xie, H.-X., Liang, X.-X., Chen, Z.-Q., Li, W.-M., Mi, C.-R., Li, M., Wu, Z.-J., Zhou, X.-M., & Du, W.-G. (2022). Ancient demographics determine the effectiveness of genetic purging in endangered lizards. Molecular Biology and Evolution, 39(1), msab359. 10.1093/molbev/msab359

Yang, L., Jin, H., Yang, Q., Poyarkov, A., Korablev, M., Rozhnov, V., Shao, J., Fu, Q., Hernandez-Blanco, J. A., Zhan, X., Yu, L., Alexandrov, D., Dai, Q., Munkhtsog, B., Du, X., Munkhtsog, B., Ma, L., Chen, W., Malykh, S., … Hu, Y. (2025). Genomic evidence for low genetic diversity but purging of strong deleterious variants in snow leopards. Genome Biology, 26(1), 94. 10.1186/s13059-025-03555-0

Yang, W., Feiner, N., Pinho, C., While, G. M., Kaliontzopoulou, A., Harris, D. J., Salvi, D., & Uller, T. (2021). Extensive introgression and mosaic genomes of Mediterranean endemic lizards. Nature Communications, 12(1), Article 1. 10.1038/s41467-021-22949-9

Yang, W., Li, C., & Mansour, S. L. (2001). Impaired motor coordination in mice that lack *punc*. Molecular and Cellular Biology, 21(17), 6031–6043. 10.1128/MCB.21.17.6031-6043.2001

Yurchenko, A. A., Recknagel, H., & Elmer, K. R. (2020). Chromosome-Level Assembly of the Common Lizard (*Zootoca vivipara*) Genome. Genome Biology and Evolution, 12(11), 1953–1960. 10.1093/gbe/evaa161

Zamora-Camacho, F. J., Reguera, S., Rubiño-Hispán, M. V., & Moreno-Rueda, G. (2015). Eliciting an immune response reduces sprint speed in a lizard. Behavioral Ecology, 26(1), 115–120. 10.1093/beheco/aru170

Zhou, X., & Stephens, M. (2012). Genome-wide efficient mixed-model analysis for association studies. Nature Genetics, 44(7), 821–824. 10.1038/ng.2310

