## Supplemental Information for "Elevated genetic load is associated with reduced bite force in insular lizard populations"

national d’Histoire naturelle CNRS, Paris, France

^5^ Department of Biology, Evolutionary Morphology of Vertebrates, Ghent University, Ghent, Belgium

^6^ Naturhistorisches Museum Bern, Bern, Switzerland

^+^ Joint senior authorship

### Supplementary Methods

#### Inbreeding depression

*Lizard performance traits*

Performance data were collected for a subset of the sampled populations in May 2024, during the reproductive season (Andò et al. 1990, 1992). In total, we captured and measured 134 male lizards from the mainland populations of Vieste (VS; 26 individuals) and Termoli (TM; 28 individuals) and the island populations of Sušac (SC; 26 individuals), Pod Kopište (PK; 27 individuals), and Pod Mrčaru (PM; 27 individuals).

For each individual, we measured snout-vent-length (SVL) and head dimensions (head width, head length, and head height) with a CD-AX/APX digital caliper (0.01 mm accuracy, Mitutoyo, USA). Individual head size was then calculated as the geometric mean of the head dimensions.

All performance tests were conducted with lizards at near-optimal temperatures (35°C, Van Damme et al. 1990), with sufficient rest (> 4 hours) between tests. To measure maximal bite force, lizards were induced to bite onto two plates of a custom-built holder of an isometric Kistler force transducer (see Herrel et al. 1999). This was repeated five times, with the maximal value recorded noted as the maximal bite force of that individual.

To measure maximal sprint speed, we chased lizards along a 2 meter custom-built race-track equipped with photocells every 25 cm (Vanhooydonck et al. 2001). This was repeated three times, with an hour of rest in between runs and the maximal speed value recorded over an interval was considered to be the maximal sprint speed of that individual.

Sperm quality (sperm count, velocity, and morphology) was assessed following the protocols developed by Kahrl et al. (2021). In short, a sperm sample of each individual was collected by gently pressing its abdomen and collecting the ejaculate in a microcapillary tube (Kahrl and Cox 2017, Martínez-Torres et al. 2019). Immediately after collection, the sample was suspended in Dulbecco’s Modified Eagle Medium and 50 µL of the suspension was added to a 1% PVA-coated well slide. A one-minute video was taken under the microscope (Reichert Biovar type 300404, Austria; 400x magnification) with an iPhone 12 (Apple, USA) on a custom-build holder (1280 x 720 resolution and 30 fps). Three sperm cells per individual were manually tracked for 40-100 frames using ImageJ (Schneider et al. 2012) to determine the average sperm velocity (µm/s) for each individual. The remaining suspension was fixed with 4% paraformaldehyde and sperm count was determined using a Bürker counting chamber (depth 0.01 mm). Lastly, the sperm suspension was dried on a slide and stained with SpermBlue™ (Microptic SL, Spain) (Kahrl and Cox 2015). Sperm head, midpiece, and tail length were measured using ImageJ (Schneider et al. 2012) from photos taken with the Carl Zeiss Axioskop 20 microscope (Germany). See Kahrl et al. (2021) for full protocol details on quantifying sperm velocity, count, and morphology.

Immunocompetence was assessed by quantifying the delayed inflammatory response to a subcutaneous injection of phytohaemaglutinin (PHA), which triggers a complex immune response involving multiple cell types, resulting in swelling (Martin et al. 2006). This test provides an indication of the proinflammatory potential of an individual (Vinkler et al. 2010). For each lizard, we measured the left and right hind footpad using a CD-AX/APX digital calliper (0.01 mm accuracy, Mitutoyo, USA), after which one foot was injected with 20µL of phosphate buffer saline (PBS) and the other foot with 20µL of a 5 mg/mL PHA/PBS solution (Sigma-Aldrich, L8754). The lizards were kept in an incubator at 32°C, and after 12 hours each footpad was measured again. The PHA response was then quantified as the change in thickness of the PHA injected footpad minus the change in the control footpad, with a bigger swelling indicating better immune condition (Mugabo et al. 2010, Huyghe et al. 2010, Baeckens and Van Damme 2018).

Both ecto- and endoparasites can influence immune condition in Podarcis lizards (Garrido and Pérez-Mellado 2014). To account for this in the statistical models, we quantified ectoparasite load by counting the number of ticks and mites present on the skin of each individual. Endoparasite infection was assessed by estimating haemogregarine intensity from blood smears: blood was collected via the medial canthus of the eye using a capillary tube, smeared onto a microscope slide, fixed in methanol, and stained with modified Giemsa solution (Sigma-Aldrich, 32884). Haemogregarine parasites were then quantified as the number of infected cells per 500 erythrocytes (Carl Zeiss Axioskop 20 microscope, Germany; 400x magnification) (Garrido and Pérez-Mellado 2014, Ferreira et al. 2023).

Spatial cognition was assessed using a spatial learning task in which individuals learned the location of a “safe” shelter (De Meester et al. 2022, Gavriilidi et al. 2026) . Trials were conducted in opaque arenas (78 × 56 × 43 cm) containing a sandy substrate and four identical clay shelters, one positioned in each corner. For each individual, one of the four shelters was randomly designated as safe, whereas the entrances of the remaining three shelters were blocked with transparent covers that were invisible to the lizard. Internal landmarks (stone structures and artificial vegetation) and external visual cues (tall wooden poles) were provided for orientation. A heating lamp (60 W) was positioned above the center of the arena, allowing the lizards to maintain their preferred body temperatures. Prior to the experiments, lizards were allowed to acclimate to the arena for 24 hours. At the start of each trial, the focal lizard was placed under a transparent cover in the center of the arena. After two minutes, a predator attack was simulated by gently tapping the base of the lizard’s tail. Chasing ceased when the lizard entered the safe shelter or when 120 s had elapsed. Following each trial, lizards remained undisturbed in the safe shelter for five minutes before being returned to their home terraria. For each trial, we recorded the number of errors (i.e. attempts to enter the unsafe shelter) and trials were considered successful if lizards entered the safe shelter first. For each individual, we also recorded the number of trials required to reach the learning criterium (i.e. four out of five consecutive successful trials; a robust learning criterium tested by an exact binomial test, p<0.05). Individuals that did not reach the learning criterium (N=40 of 125, two individuals were excluded), were assigned a maximum penalty value. Lizards were tested three times per day, with minimally one hour in between trials, for six consecutive days (resulting in 18 trials in total). All spatial learning variables (average N° of errors, N° of successful trials, and N° of trials to reach the learning criterion) were highly correlated (ρ=0.50-0.82; p<0.005), and N° of trials to reach the learning criterion was selected as the spatial learning measure for all subsequent analyses.

### References

Alexander, D. H., Novembre, J. and Lange, K. 2009. Fast model-based estimation of ancestry in unrelated individuals. - Genome Res. 19: 1655.

Andò, S., Panno, M. L., Ciarcia, G., Imbrogno, E., Buffone, M., Beraldi, E., Sisci, D., Angelini, F. and Botte, V. 1990. Plasma sex hormone concentrations during the reproductive cycle in the male lizard, Podarcis s. sicula. - Reproduction 90: 353–360.

Andò, S., Ciarcia, G., Panno, M. L., Imbrogno, E., Tarantino, G., Buffone, M., Beraldi, E., Angelini, F. and Botte, V. 1992. Sex steroids levels in the plasma and testis during the reproductive cycle of lizard *Podarcis s. sicula raf*. - Gen. Comp. Endocrinol. 85: 1–7.

Baeckens, S. and Van Damme, R. 2018. Immunocompetence and parasite infestation in a melanistic and normally-coloured population of the lacertid lizard, Podarcis siculus. - Amphib.-Reptil. in press.

De Meester, G., Van Linden, L., Torfs, J., Pafilis, P., Šunje, E., Steenssens, D., Zulčić, T., Sassalos, A. and Van Damme, R. 2022. Learning with lacertids: Studying the link between ecology and cognition within a comparative framework. - Evolution 76: 2531–2552.

Ferreira, A. I., Damas-Moreira, I., Marshall, K. L. A., Perera, A. and Harris, D. J. 2023. What Influences the Prevalence and Intensity of Haemoparasites and Ectoparasites in an Insular Lizard? - Animals 13: 723.

Garrido, M. and Pérez-Mellado, V. 2014. Immune responses vary with parasite burden in an insular lizard. - Evolutionary Ecology Research 16: 593–603.

Gavriilidi, I., Baeckens, S., Pafilis, P. and Van Damme, R. 2026. Creatures of habit: similar but more repeatable behaviours in insular compared with mainland lizards. - Anim. Behav. 233: 123493.

Herrel, A., Spithoven, L., Van Damme, R. and DE Vree, F. 1999. Sexual dimorphism of head size in Gallotia galloti: testing the niche divergence hypothesis by functional analyses. - Funct. Ecol. 13: 289–297.

Huyghe, K., Van Oystaeyen, A., Pasmans, F., Tadić, Z., Vanhooydonck, B. and Van Damme, R. 2010. Seasonal changes in parasite load and a cellular immune response in a colour polymorphic lizard. - Oecologia 163: 867–874.

Kahrl, A. F. and Cox, R. M. 2015. Diet affects ejaculate traits in a lizard with condition-dependent fertilization success. - Behav. Ecol. 26: 1502–1511.

Kahrl, A. F. and Cox, R. M. 2017. Consistent Differences in Sperm Morphology and Testis Size between Native and Introduced Populations of Three Anolis Lizard Species. - J. Herpetol. 51: 532–537.

Kahrl, A. F., Kustra, M. C., Reedy, A. M., Bhave, R. S., Seears, H. A., Warner, D. A. and Cox, R. M. 2021. Selection on Sperm Count, but Not on Sperm Morphology or Velocity, in a Wild Population of Anolis Lizards. - Cells 10: 2369.

Li, H., Handsaker, B., Wysoker, A., Fennell, T., Ruan, J., Homer, N., Marth, G., Abecasis, G., Durbin, R., and 1000 Genome Project Data Processing Subgroup 2009. The Sequence Alignment/Map format and SAMtools. - Bioinformatics 25: 2078–2079.

Martin, L. B., Han, P., Lewittes, J., Kuhlman, J. R., Klasing, K. C. and Wikelski, M. 2006. Phytohemagglutinin-induced skin swelling in birds: histological support for a classic immunoecological technique. - Funct. Ecol. 20: 290–299.

Martínez-Torres, M., Sánchez-Rivera, U. Á., Cruz-Cano, N. B., Castro-Camacho, Y. J., Luis, J. and Medrano, A. 2019. A non-invasive method for semen collection and evaluation in small and median size lizards. - Reprod. Domest. Anim. 54: 54–58.

Mugabo, M., Marquis, O., Perret, S. and Le Galliard, J. F. 2010. Immediate and delayed life history effects caused by food deprivation early in life in a short-lived lizard. - J. Evol. Biol. 23: 1886–1898.

Ramírez, F., Ryan, D. P., Grüning, B., Bhardwaj, V., Kilpert, F., Richter, A. S., Heyne, S., Dündar, F. and Manke, T. 2016. deepTools2: a next generation web server for deep-sequencing data analysis. - Nucleic Acids Res. 44: W160–W165.

Schneider, C. A., Rasband, W. S. and Eliceiri, K. W. 2012. NIH Image to ImageJ: 25 years of image analysis. - Nat. Methods 9: 671–675.

Van Damme, R., Bauwens, D., Castilla, A. M. and Verheyen, R. F. 1990. Comparative thermal ecology of the sympatric lizards *Podarcis tiliguerta* and *Podarcis sicula*. - Acta Oecologica Montrouge 11: 503–512.

Vanhooydonck, B., Van Damme, R. and Aerts, P. 2001. Speed and stamina trade-off in Lacertid lizards. - Evolution 55: 1040–1048.

Vinkler, M., Bainová, H. and Albrecht, T. 2010. Functional analysis of the skin-swelling response to phytohaemagglutinin. - Funct. Ecol. 24: 1081–1086.

**Table S1:** Overview of sampled *Podarcis siculus* populations, including population ID, locality, country, coordinates of the sampling location, sample sizes per sampling year. Previously published genomic data are referenced.

| **Population ID** | **Locality** | **Country** | **Coordinates** | **Population type** | **Sampling year** | **N** | **Genomic data referenced in** | **Total N** |
| --- | --- | --- | --- | --- | --- | --- | --- | --- |
| VS | Vieste | Italy | 41°51'45.0"N 16°09'54.0"E | mainland | 2022 | 2 | This paper | 38 |
|  |  |  |  |  | 2023 | 10 |  |  |
|  |  |  |  |  | 2024 | 26 |  |  |
| RG | Rodi Garganico | Italy | 41°55'31.0"N 15°52'1.0"E | mainland | 2022 | 8 | This paper | 16 |
|  |  |  |  |  | 2023 | 8 |  |  |
| TM | Termoli | Italy | 41°58'42.0"N 14°56'42.0"E | mainland | 2022 | 3 | This paper | 40 |
|  |  |  |  |  | 2023 | 9 |  |  |
|  |  |  |  |  | 2024 | 28 |  |  |
| PM | Pod Mrčaru | Croatia | 42°46'47.8"N 16°46'29.5"E | island | 2023 | 14 | Van Linden et al. (2025) Oecologia | 46 |
|  |  |  |  |  | 2024 | 32 | This paper |  |
| PK | Pod Kopište | Croatia | 42°45'47.4"N 16°43'27.0"E | island | 2023 | 14 | Van Linden et al. (2025) Oecologia | 46 |
|  |  |  |  |  | 2024 | 32 | This paper |  |
| KP | Kopište | Croatia | 42°45'2.8"N 16°43'7.8"E | island | 2023 | 13 | Van Linden et al. (2025) Oecologia | 13 |
| SC | Sušac | Croatia | 42°45'38.3"N 16°31'45.5"E | island | 2023 | 12 | Van Linden et al. (2025) Oecologia | 43 |
|  |  |  |  |  | 2024 | 31 | This paper |  |
| PJ | Pijavica | Croatia | 43°28'32.2"N 16°11'26.4"E | island | 2016 | 3 | This paper | 3 |
| MP | Mala Palagruža | Croatia | 42°23'18.1"N 16°16'18.5"E | island | 2016 | 3 | This paper | 3 |
| BJ | Bijelac | Croatia | 42°45'31.4"N 16°40'45.1"E | island | 2016 | 5 | Van Linden et al. (2025) Oecologia | 5 |
| PL | Paklenica | Croatia | 44°14'03.7"N 15°32'09.5"E | mainland | 2004 | 3 | This paper | 3 |
| SP | Split | Croatia | 43°30'50.4"N 16°26'37.7"E | mainland | 2004 | 2 | This paper | 2 |
| TR | Trogir | Croatia | 43°31'12.9"N 16°15'30.1"E | mainland | 2024 | 5 | This paper | 5 |

**Table S2:** Overview of all samples, including locality, type, sample ID, mean genome-wide coverage, number (in millions, M) and percentage (%) of reads. Mean genome-wide coverage and number and percentage of reads mapped for all samples in the final dataset obtained with deepTools2.0 (Ramírez et al. 2016) and SAMtools ‘flagstat’ (Li et al. 2009). Previously published genomic data are referenced.

| **Locality** | **Population type** | **Sample ID** | **Mean coverage (x)** | **M reads mapped** | **% reads mapped** | **Referenced in** |
| --- | --- | --- | --- | --- | --- | --- |
| Bijelac | island | BJ04 | 15.56 | 164.5 | 99.60% | Van Linden et al. (2025) Oecologia |
| Bijelac | island | BJ06 | 15.59 | 164.9 | 99.60% | Van Linden et al. (2025) Oecologia |
| Bijelac | island | BJ09 | 15.65 | 165.5 | 99.60% | Van Linden et al. (2025) Oecologia |
| Bijelac | island | BJ16 | 15.57 | 164.7 | 99.60% | Van Linden et al. (2025) Oecologia |
| Bijelac | island | BJ18 | 15.61 | 165.1 | 99.60% | Van Linden et al. (2025) Oecologia |
| Kopište | island | KP02 | 15.52 | 164.1 | 99.40% | Van Linden et al. (2025) Oecologia |
| Kopište | island | KP03 | 15.55 | 164.4 | 99.60% | Van Linden et al. (2025) Oecologia |
| Kopište | island | KP04 | 15.6 | 165 | 99.50% | Van Linden et al. (2025) Oecologia |
| Kopište | island | KP05 | 14.66 | 155 | 99.50% | Van Linden et al. (2025) Oecologia |
| Kopište | island | KP06 | 15.52 | 164.1 | 99.30% | Van Linden et al. (2025) Oecologia |
| Kopište | island | KP07 | 15.52 | 164.2 | 99.50% | Van Linden et al. (2025) Oecologia |
| Kopište | island | KP08 | 15.5 | 163.9 | 99.30% | Van Linden et al. (2025) Oecologia |
| Kopište | island | KP11 | 15.55 | 164.4 | 99.60% | Van Linden et al. (2025) Oecologia |
| Kopište | island | KP12 | 15.56 | 164.5 | 99.50% | Van Linden et al. (2025) Oecologia |
| Kopište | island | KP13 | 15.52 | 164.1 | 99.40% | Van Linden et al. (2025) Oecologia |
| Kopište | island | KP14 | 15.54 | 164.3 | 99.50% | Van Linden et al. (2025) Oecologia |
| Kopište | island | KP15 | 15.55 | 164.5 | 99.60% | Van Linden et al. (2025) Oecologia |
| Kopište | island | KP16 | 15.52 | 164.1 | 99.40% | Van Linden et al. (2025) Oecologia |
| Mala Palagruža | island | MP03 | 15.62 | 165.1 | 99.60% | This paper |
| Mala Palagruža | island | MP07 | 15.61 | 165.1 | 99.64% | This paper |
| Mala Palagruža | island | MP13 | 15.64 | 165.4 | 99.56% | This paper |
| Pijavica | island | PJ14 | 15.69 | 165.9 | 99.65% | This paper |
| Pijavica | island | PJ17 | 15.65 | 165.5 | 99.62% | This paper |
| Pijavica | island | PJ19 | 15.64 | 165.3 | 99.65% | This paper |
| Pod Kopište | island | 24PK01 | 15.57 | 164.6 | 99.54% | This paper |
| Pod Kopište | island | 24PK02 | 15.57 | 164.6 | 99.69% | This paper |
| Pod Kopište | island | 24PK03 | 15.69 | 165.9 | 99.27% | This paper |
| Pod Kopište | island | 24PK04 | 15.66 | 165.6 | 99.66% | This paper |
| Pod Kopište | island | 24PK05 | 15.82 | 167.2 | 99.64% | This paper |
| Pod Kopište | island | 24PK06 | 15.87 | 167.8 | 99.75% | This paper |
| Pod Kopište | island | 24PK07 | 15.61 | 165.1 | 99.73% | This paper |
| Pod Kopište | island | 24PK08 | 15.86 | 167.7 | 99.73% | This paper |
| Pod Kopište | island | 24PK09 | 15.69 | 165.9 | 99.72% | This paper |
| Pod Kopište | island | 24PK10 | 15.62 | 165.2 | 99.70% | This paper |
| Pod Kopište | island | 24PK11 | 15.87 | 167.8 | 99.64% | This paper |
| Pod Kopište | island | 24PK12 | 16.32 | 172.6 | 98.59% | This paper |
| Pod Kopište | island | 24PK13 | 15.59 | 164.8 | 99.75% | This paper |
| Pod Kopište | island | 24PK14 | 13.75 | 145.4 | 99.47% | This paper |
| Pod Kopište | island | 24PK15 | 15.63 | 165.3 | 99.70% | This paper |
| Pod Kopište | island | 24PK16 | 15.53 | 164.2 | 99.49% | This paper |
| Pod Kopište | island | 24PK17 | 15.57 | 164.6 | 99.69% | This paper |
| Pod Kopište | island | 24PK18 | 15.6 | 165 | 99.74% | This paper |
| Pod Kopište | island | 24PK19 | 15.57 | 164.7 | 99.67% | This paper |
| Pod Kopište | island | 24PK20 | 15.58 | 164.7 | 99.76% | This paper |
| Pod Kopište | island | 24PK21 | 13.22 | 139.8 | 99.83% | This paper |
| Pod Kopište | island | 24PK22 | 15.78 | 166.8 | 99.73% | This paper |
| Pod Kopište | island | 24PK23 | 15.47 | 163.6 | 99.06% | This paper |
| Pod Kopište | island | 24PK24 | 15.61 | 165.1 | 99.73% | This paper |
| Pod Kopište | island | 24PK25 | 15.72 | 166.2 | 99.60% | This paper |
| Pod Kopište | island | 24PK26 | 15.67 | 165.7 | 99.72% | This paper |
| Pod Kopište | island | 24PK27 | 15.72 | 166.2 | 99.73% | This paper |
| Pod Kopište | island | 24PK28 | 16.1 | 170.3 | 99.66% | This paper |
| Pod Kopište | island | 24PK29 | 15.83 | 167.4 | 99.59% | This paper |
| Pod Kopište | island | 24PK30 | 15.73 | 166.4 | 99.68% | This paper |
| Pod Kopište | island | 24PK31 | 15.83 | 167.4 | 99.33% | This paper |
| Pod Kopište | island | 24PK32 | 15.85 | 167.6 | 99.75% | This paper |
| Pod Kopište | island | PK01 | 15.62 | 165.1 | 99.50% | Van Linden et al. (2025) Oecologia |
| Pod Kopište | island | PK02 | 15.6 | 164.9 | 99.50% | Van Linden et al. (2025) Oecologia |
| Pod Kopište | island | PK03 | 15.63 | 165.3 | 99.60% | Van Linden et al. (2025) Oecologia |
| Pod Kopište | island | PK04 | 15.58 | 164.7 | 99.60% | Van Linden et al. (2025) Oecologia |
| Pod Kopište | island | PK05 | 15.56 | 164.5 | 99.40% | Van Linden et al. (2025) Oecologia |
| Pod Kopište | island | PK06 | 13.62 | 144 | 99.40% | Van Linden et al. (2025) Oecologia |
| Pod Kopište | island | PK07 | 15.54 | 164.3 | 99.40% | Van Linden et al. (2025) Oecologia |
| Pod Kopište | island | PK08 | 15.54 | 164.3 | 99.40% | Van Linden et al. (2025) Oecologia |
| Pod Kopište | island | PK09 | 15.55 | 164.4 | 99.50% | Van Linden et al. (2025) Oecologia |
| Pod Kopište | island | PK121 | 15.63 | 165.2 | 99.60% | Van Linden et al. (2025) Oecologia |
| Pod Kopište | island | PK122 | 15.62 | 165.2 | 99.60% | Van Linden et al. (2025) Oecologia |
| Pod Kopište | island | PK123 | 15.71 | 166.1 | 99.70% | Van Linden et al. (2025) Oecologia |
| Pod Kopište | island | PK124 | 15.91 | 168.2 | 99.70% | Van Linden et al. (2025) Oecologia |
| Pod Kopište | island | PK125 | 15.69 | 166 | 99.60% | Van Linden et al. (2025) Oecologia |
| Pod Mrčaru | island | 24PM01 | 15.56 | 164.5 | 99.73% | This paper |
| Pod Mrčaru | island | 24PM02 | 15.56 | 164.5 | 99.78% | This paper |
| Pod Mrčaru | island | 24PM03 | 15.55 | 164.5 | 99.72% | This paper |
| Pod Mrčaru | island | 24PM04 | 15.49 | 163.8 | 99.52% | This paper |
| Pod Mrčaru | island | 24PM05 | 15.57 | 164.6 | 99.78% | This paper |
| Pod Mrčaru | island | 24PM06 | 15.54 | 164.3 | 99.67% | This paper |
| Pod Mrčaru | island | 24PM07 | 15.5 | 163.9 | 99.38% | This paper |
| Pod Mrčaru | island | 24PM08 | 15.54 | 164.3 | 99.71% | This paper |
| Pod Mrčaru | island | 24PM09 | 15.55 | 164.5 | 99.44% | This paper |
| Pod Mrčaru | island | 24PM10 | 15.56 | 164.5 | 99.41% | This paper |
| Pod Mrčaru | island | 24PM11 | 15.51 | 164.1 | 99.44% | This paper |
| Pod Mrčaru | island | 24PM12 | 15.5 | 164 | 99.56% | This paper |
| Pod Mrčaru | island | 24PM13 | 15.54 | 164.3 | 99.70% | This paper |
| Pod Mrčaru | island | 24PM14 | 15.55 | 164.4 | 99.73% | This paper |
| Pod Mrčaru | island | 24PM15 | 15.52 | 164.1 | 99.53% | This paper |
| Pod Mrčaru | island | 24PM16 | 15.56 | 164.5 | 99.74% | This paper |
| Pod Mrčaru | island | 24PM17 | 15.5 | 163.9 | 99.54% | This paper |
| Pod Mrčaru | island | 24PM18 | 15.56 | 164.5 | 99.69% | This paper |
| Pod Mrčaru | island | 24PM19 | 15.52 | 164.1 | 99.64% | This paper |
| Pod Mrčaru | island | 24PM20 | 15.56 | 164.5 | 99.71% | This paper |
| Pod Mrčaru | island | 24PM21 | 15.53 | 164.2 | 99.70% | This paper |
| Pod Mrčaru | island | 24PM22 | 15.56 | 164.6 | 99.64% | This paper |
| Pod Mrčaru | island | 24PM23 | 15.51 | 164 | 99.62% | This paper |
| Pod Mrčaru | island | 24PM24 | 15.54 | 164.4 | 99.73% | This paper |
| Pod Mrčaru | island | 24PM25 | 15.56 | 164.5 | 99.75% | This paper |
| Pod Mrčaru | island | 24PM26 | 15.58 | 164.7 | 99.73% | This paper |
| Pod Mrčaru | island | 24PM27 | 15.68 | 165.8 | 99.73% | This paper |
| Pod Mrčaru | island | 24PM28 | 15.61 | 165.1 | 99.76% | This paper |
| Pod Mrčaru | island | 24PM29 | 15.68 | 165.8 | 99.73% | This paper |
| Pod Mrčaru | island | 24PM30 | 15.96 | 168.8 | 99.80% | This paper |
| Pod Mrčaru | island | 24PM31 | 15.72 | 166.2 | 99.77% | This paper |
| Pod Mrčaru | island | 24PM32 | 15.61 | 165.1 | 99.76% | This paper |
| Pod Mrčaru | island | PM01 | 15.54 | 164.3 | 99.40% | Van Linden et al. (2025) Oecologia |
| Pod Mrčaru | island | PM03 | 15.55 | 164.4 | 99.50% | Van Linden et al. (2025) Oecologia |
| Pod Mrčaru | island | PM04 | 15.59 | 164.8 | 99.40% | Van Linden et al. (2025) Oecologia |
| Pod Mrčaru | island | PM05 | 15.46 | 163.5 | 99.00% | Van Linden et al. (2025) Oecologia |
| Pod Mrčaru | island | PM06 | 15.66 | 165.6 | 99.60% | Van Linden et al. (2025) Oecologia |
| Pod Mrčaru | island | PM07 | 15.52 | 164.1 | 99.20% | Van Linden et al. (2025) Oecologia |
| Pod Mrčaru | island | PM08 | 14.32 | 151.4 | 99.60% | Van Linden et al. (2025) Oecologia |
| Pod Mrčaru | island | PM09 | 15.64 | 165.4 | 99.50% | Van Linden et al. (2025) Oecologia |
| Pod Mrčaru | island | PM10 | 15.52 | 164.1 | 99.30% | Van Linden et al. (2025) Oecologia |
| Pod Mrčaru | island | PM13 | 15.56 | 164.5 | 99.50% | Van Linden et al. (2025) Oecologia |
| Pod Mrčaru | island | PM14 | 15.59 | 164.9 | 99.60% | Van Linden et al. (2025) Oecologia |
| Pod Mrčaru | island | PM15 | 15.6 | 164.9 | 99.50% | Van Linden et al. (2025) Oecologia |
| Pod Mrčaru | island | PM16 | 13.72 | 145 | 99.00% | Van Linden et al. (2025) Oecologia |
| Pod Mrčaru | island | PM17 | 15.63 | 165.2 | 99.50% | Van Linden et al. (2025) Oecologia |
| Sušac | island | 24SC01 | 14.11 | 149.2 | 98.93% | This paper |
| Sušac | island | 24SC02 | 15.59 | 164.8 | 99.64% | This paper |
| Sušac | island | 24SC03 | 14.25 | 150.7 | 99.73% | This paper |
| Sušac | island | 24SC04 | 15.49 | 163.8 | 99.48% | This paper |
| Sušac | island | 24SC05 | 15.51 | 164 | 99.32% | This paper |
| Sušac | island | 24SC06 | 15.71 | 166.1 | 99.38% | This paper |
| Sušac | island | 24SC07 | 14.27 | 150.9 | 99.65% | This paper |
| Sušac | island | 24SC08 | 15.59 | 164.9 | 99.58% | This paper |
| Sušac | island | 24SC09 | 15.56 | 164.5 | 99.65% | This paper |
| Sušac | island | 24SC10 | 14.68 | 155.3 | 99.59% | This paper |
| Sušac | island | 24SC11 | 15.57 | 164.7 | 99.64% | This paper |
| Sušac | island | 24SC12 | 14.87 | 157.2 | 99.61% | This paper |
| Sušac | island | 24SC13 | 15.62 | 165.2 | 99.36% | This paper |
| Sušac | island | 24SC14 | 15.56 | 165.6 | 99.24% | This paper |
| Sušac | island | 24SC15 | 15.42 | 163 | 99.69% | This paper |
| Sušac | island | 24SC16 | 15.57 | 164.7 | 99.60% | This paper |
| Sušac | island | 24SC17 | 13.03 | 137.8 | 99.65% | This paper |
| Sušac | island | 24SC18 | 15.57 | 165.7 | 99.72% | This paper |
| Sušac | island | 24SC19 | 15.56 | 164.5 | 99.63% | This paper |
| Sušac | island | 24SC20 | 15.66 | 165.5 | 99.68% | This paper |
| Sušac | island | 24SC21 | 15.56 | 164.5 | 99.64% | This paper |
| Sušac | island | 24SC22 | 15.57 | 164.6 | 99.68% | This paper |
| Sušac | island | 24SC23 | 16.39 | 173.3 | 99.78% | This paper |
| Sušac | island | 24SC24 | 15.58 | 164.8 | 99.50% | This paper |
| Sušac | island | 24SC25 | 15.68 | 165.9 | 99.77% | This paper |
| Sušac | island | 24SC26 | 15.75 | 166.6 | 99.70% | This paper |
| Sušac | island | 24SC27 | 15.74 | 166.5 | 99.60% | This paper |
| Sušac | island | 24SC28 | 15.74 | 166.5 | 99.68% | This paper |
| Sušac | island | 24SC29 | 15.69 | 165.9 | 99.63% | This paper |
| Sušac | island | 24SC30 | 15.85 | 167.6 | 99.69% | This paper |
| Sušac | island | 24SC31 | 15.85 | 167.6 | 99.66% | This paper |
| Sušac | island | SC01 | 15.53 | 164.3 | 99.50% | Van Linden et al. (2025) Oecologia |
| Sušac | island | SC02 | 15.56 | 164.5 | 99.60% | Van Linden et al. (2025) Oecologia |
| Sušac | island | SC03 | 15.5 | 163.8 | 99.30% | Van Linden et al. (2025) Oecologia |
| Sušac | island | SC04 | 15.51 | 164 | 99.40% | Van Linden et al. (2025) Oecologia |
| Sušac | island | SC05 | 15.52 | 164.2 | 99.50% | Van Linden et al. (2025) Oecologia |
| Sušac | island | SC06 | 15.46 | 163.4 | 98.90% | Van Linden et al. (2025) Oecologia |
| Sušac | island | SC07 | 15.56 | 164.5 | 99.60% | Van Linden et al. (2025) Oecologia |
| Sušac | island | SC08 | 15.54 | 164.3 | 99.50% | Van Linden et al. (2025) Oecologia |
| Sušac | island | SC09 | 15.52 | 164.1 | 99.30% | Van Linden et al. (2025) Oecologia |
| Sušac | island | SC10 | 15.4 | 162.9 | 98.80% | Van Linden et al. (2025) Oecologia |
| Sušac | island | SC11 | 15.52 | 164.2 | 99.40% | Van Linden et al. (2025) Oecologia |
| Sušac | island | SC12 | 15.52 | 164.2 | 99.40% | Van Linden et al. (2025) Oecologia |
| Paklenica | mainland | PL01 | 15.56 | 164.5 | 99.85% | This paper |
| Paklenica | mainland | PL02 | 14.03 | 148.4 | 99.84% | This paper |
| Paklenica | mainland | PL03 | 15.58 | 164.8 | 99.86% | This paper |
| Rodi Garganico | mainland | CV61 | 19.47 | 205.8 | 99.67% | This paper |
| Rodi Garganico | mainland | CV68 | 19.52 | 206.5 | 99.68% | This paper |
| Rodi Garganico | mainland | FV44 | 19.42 | 205.3 | 99.62% | This paper |
| Rodi Garganico | mainland | FV46 | 19.5 | 206.2 | 99.69% | This paper |
| Rodi Garganico | mainland | RG01 | 15.55 | 164.4 | 99.41% | This paper |
| Rodi Garganico | mainland | RG02 | 15.58 | 164.7 | 99.47% | This paper |
| Rodi Garganico | mainland | RG03 | 15.59 | 164.9 | 99.59% | This paper |
| Rodi Garganico | mainland | RG04 | 15.6 | 164.9 | 99.62% | This paper |
| Rodi Garganico | mainland | RG05 | 15.42 | 163.1 | 98.73% | This paper |
| Rodi Garganico | mainland | RG06 | 14.65 | 154.9 | 99.36% | This paper |
| Rodi Garganico | mainland | RG07 | 13.9 | 146.9 | 99.65% | This paper |
| Rodi Garganico | mainland | RG08 | 15.59 | 164.8 | 99.61% | This paper |
| Rodi Garganico | mainland | RG21 | 19.46 | 205.8 | 99.70% | This paper |
| Rodi Garganico | mainland | RG26 | 15.63 | 165.3 | 99.11% | This paper |
| Rodi Garganico | mainland | RG28 | 15.75 | 166.6 | 99.70% | This paper |
| Rodi Garganico | mainland | RG32 | 19.63 | 207.6 | 99.70% | This paper |
| Split | mainland | SP19 | 15.51 | 164 | 99.85% | This paper |
| Split | mainland | SP33 | 14.09 | 149 | 99.70% | This paper |
| Termoli | mainland | 24TM01 | 15.59 | 164.9 | 99.75% | This paper |
| Termoli | mainland | 24TM02 | 15.77 | 166.7 | 99.74% | This paper |
| Termoli | mainland | 24TM03 | 15.58 | 164.7 | 99.75% | This paper |
| Termoli | mainland | 24TM04 | 15.62 | 165.1 | 99.65% | This paper |
| Termoli | mainland | 24TM05 | 15.63 | 165.3 | 99.76% | This paper |
| Termoli | mainland | 24TM06 | 15.61 | 165.1 | 99.77% | This paper |
| Termoli | mainland | 24TM07 | 15.69 | 165.9 | 99.75% | This paper |
| Termoli | mainland | 24TM08 | 15.58 | 164.7 | 99.59% | This paper |
| Termoli | mainland | 24TM09 | 15.6 | 164.9 | 99.70% | This paper |
| Termoli | mainland | 24TM10 | 15.64 | 165.4 | 99.75% | This paper |
| Termoli | mainland | 24TM11 | 15.61 | 165 | 99.73% | This paper |
| Termoli | mainland | 24TM12 | 15.69 | 165.9 | 99.76% | This paper |
| Termoli | mainland | 24TM13 | 15.48 | 163.7 | 99.25% | This paper |
| Termoli | mainland | 24TM14 | 15.56 | 164.5 | 99.41% | This paper |
| Termoli | mainland | 24TM15 | 15.65 | 165.5 | 99.77% | This paper |
| Termoli | mainland | 24TM16 | 15.68 | 165.8 | 99.73% | This paper |
| Termoli | mainland | 24TM17 | 15.83 | 167.4 | 99.80% | This paper |
| Termoli | mainland | 24TM18 | 15.63 | 165.3 | 99.73% | This paper |
| Termoli | mainland | 24TM19 | 15.54 | 164.3 | 99.39% | This paper |
| Termoli | mainland | 24TM20 | 16.05 | 169.7 | 99.72% | This paper |
| Termoli | mainland | 24TM21 | 15.61 | 165.1 | 99.75% | This paper |
| Termoli | mainland | 24TM22 | 15.65 | 165.5 | 99.75% | This paper |
| Termoli | mainland | 24TM23 | 15.6 | 165 | 99.61% | This paper |
| Termoli | mainland | 24TM24 | 15.75 | 166.6 | 99.69% | This paper |
| Termoli | mainland | 24TM25 | 15.64 | 165.4 | 99.70% | This paper |
| Termoli | mainland | 24TM26 | 15.7 | 166 | 99.55% | This paper |
| Termoli | mainland | 24TM27 | 15.79 | 167 | 99.70% | This paper |
| Termoli | mainland | 24TM28 | 15.81 | 167.2 | 99.75% | This paper |
| Termoli | mainland | TM01 | 15.41 | 163 | 98.24% | This paper |
| Termoli | mainland | TM02 | 15.66 | 165.6 | 99.65% | This paper |
| Termoli | mainland | TM03 | 15.58 | 164.8 | 99.42% | This paper |
| Termoli | mainland | TM04 | 15.62 | 165.2 | 99.61% | This paper |
| Termoli | mainland | TM05 | 15.58 | 164.8 | 99.45% | This paper |
| Termoli | mainland | TM06 | 15.64 | 165.4 | 99.63% | This paper |
| Termoli | mainland | TM07 | 15.64 | 165.3 | 99.60% | This paper |
| Termoli | mainland | TM08 | 14.99 | 158.5 | 97.29% | This paper |
| Termoli | mainland | TM09 | 14.73 | 155.7 | 94.15% | This paper |
| Termoli | mainland | TM84 | 19.49 | 206.1 | 99.68% | This paper |
| Termoli | mainland | TM85 | 15.71 | 166.1 | 99.68% | This paper |
| Termoli | mainland | TM96 | 19.32 | 204.3 | 98.66% | This paper |
| Trogir | mainland | TR01 | 15.6 | 164.9 | 99.74% | This paper |
| Trogir | mainland | TR04 | 14.27 | 151 | 99.76% | This paper |
| Trogir | mainland | TR07 | 15.63 | 165.2 | 99.74% | This paper |
| Trogir | mainland | TR08 | 15.57 | 164.7 | 99.75% | This paper |
| Trogir | mainland | TR09 | 15.62 | 165.2 | 99.76% | This paper |
| Vieste | mainland | 24VS01 | 15.91 | 168.2 | 99.74% | This paper |
| Vieste | mainland | 24VS02 | 15.63 | 165.3 | 99.71% | This paper |
| Vieste | mainland | 24VS03 | 15.6 | 164.9 | 99.76% | This paper |
| Vieste | mainland | 24VS04 | 15.59 | 164.8 | 99.74% | This paper |
| Vieste | mainland | 24VS05 | 15.61 | 165 | 99.71% | This paper |
| Vieste | mainland | 24VS06 | 15.81 | 167.1 | 99.77% | This paper |
| Vieste | mainland | 24VS07 | 15.58 | 164.7 | 99.76% | This paper |
| Vieste | mainland | 24VS08 | 15.65 | 165.5 | 99.74% | This paper |
| Vieste | mainland | 24VS09 | 15.81 | 167.2 | 99.74% | This paper |
| Vieste | mainland | 24VS10 | 15.62 | 165.1 | 99.62% | This paper |
| Vieste | mainland | 24VS11 | 15.69 | 165.9 | 99.74% | This paper |
| Vieste | mainland | 24VS12 | 15.67 | 165.7 | 99.75% | This paper |
| Vieste | mainland | 24VS13 | 15.75 | 166.5 | 99.66% | This paper |
| Vieste | mainland | 24VS14 | 15.59 | 164.8 | 99.72% | This paper |
| Vieste | mainland | 24VS15 | 15.55 | 164.5 | 99.52% | This paper |
| Vieste | mainland | 24VS16 | 15.62 | 165.2 | 99.73% | This paper |
| Vieste | mainland | 24VS17 | 15.61 | 165.1 | 99.76% | This paper |
| Vieste | mainland | 24VS18 | 15.64 | 165.3 | 99.71% | This paper |
| Vieste | mainland | 24VS19 | 15.62 | 165.1 | 99.57% | This paper |
| Vieste | mainland | 24VS20 | 15.64 | 165.3 | 99.77% | This paper |
| Vieste | mainland | 24VS21 | 15.57 | 164.6 | 99.71% | This paper |
| Vieste | mainland | 24VS22 | 15.51 | 164 | 99.04% | This paper |
| Vieste | mainland | 24VS23 | 15.7 | 166 | 99.76% | This paper |
| Vieste | mainland | 24VS24 | 15.91 | 168.3 | 99.74% | This paper |
| Vieste | mainland | 24VS25 | 15.5 | 163.9 | 99.18% | This paper |
| Vieste | mainland | 24VS26 | 15.68 | 165.8 | 99.77% | This paper |
| Vieste | mainland | VS01 | 15.83 | 167.4 | 99.52% | This paper |
| Vieste | mainland | VS02 | 15.65 | 165.5 | 99.65% | This paper |
| Vieste | mainland | VS03 | 15.56 | 164.5 | 99.32% | This paper |
| Vieste | mainland | VS04 | 15.61 | 165.1 | 99.59% | This paper |
| Vieste | mainland | VS05 | 15.61 | 165.1 | 99.48% | This paper |
| Vieste | mainland | VS06 | 14.77 | 156.2 | 99.63% | This paper |
| Vieste | mainland | VS07 | 15.62 | 165.2 | 99.53% | This paper |
| Vieste | mainland | VS08 | 15.64 | 165.4 | 99.54% | This paper |
| Vieste | mainland | VS09 | 15.66 | 165.6 | 99.57% | This paper |
| Vieste | mainland | VS10 | 15.67 | 165.7 | 99.65% | This paper |
| Vieste | mainland | VS11 | 19.45 | 205.6 | 99.67% | This paper |
| Vieste | mainland | VS12 | 19.83 | 209.8 | 99.75% | This paper |

### Supplementary Results

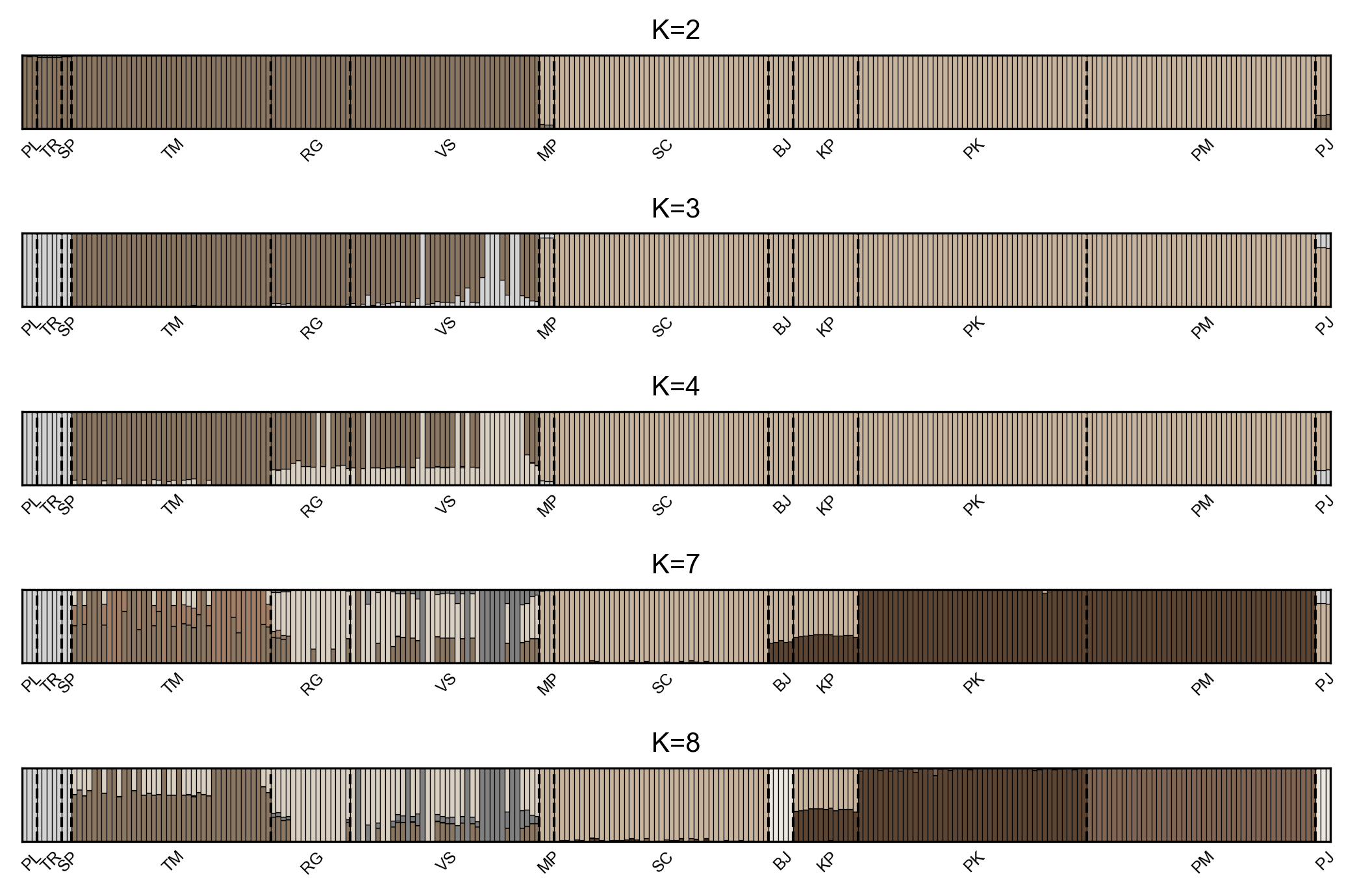

**Figure S1:** ADMIXTURE plots showing the five K-values with the lowest cross-validation (CV) error, with K = 2 being the lowest (see Table S3 for all CV errors). In each plot, different shades indicate distinct genetic clusters. Each bar represents a single *P. siculus* individual, with dashed lines marking population boundaries.

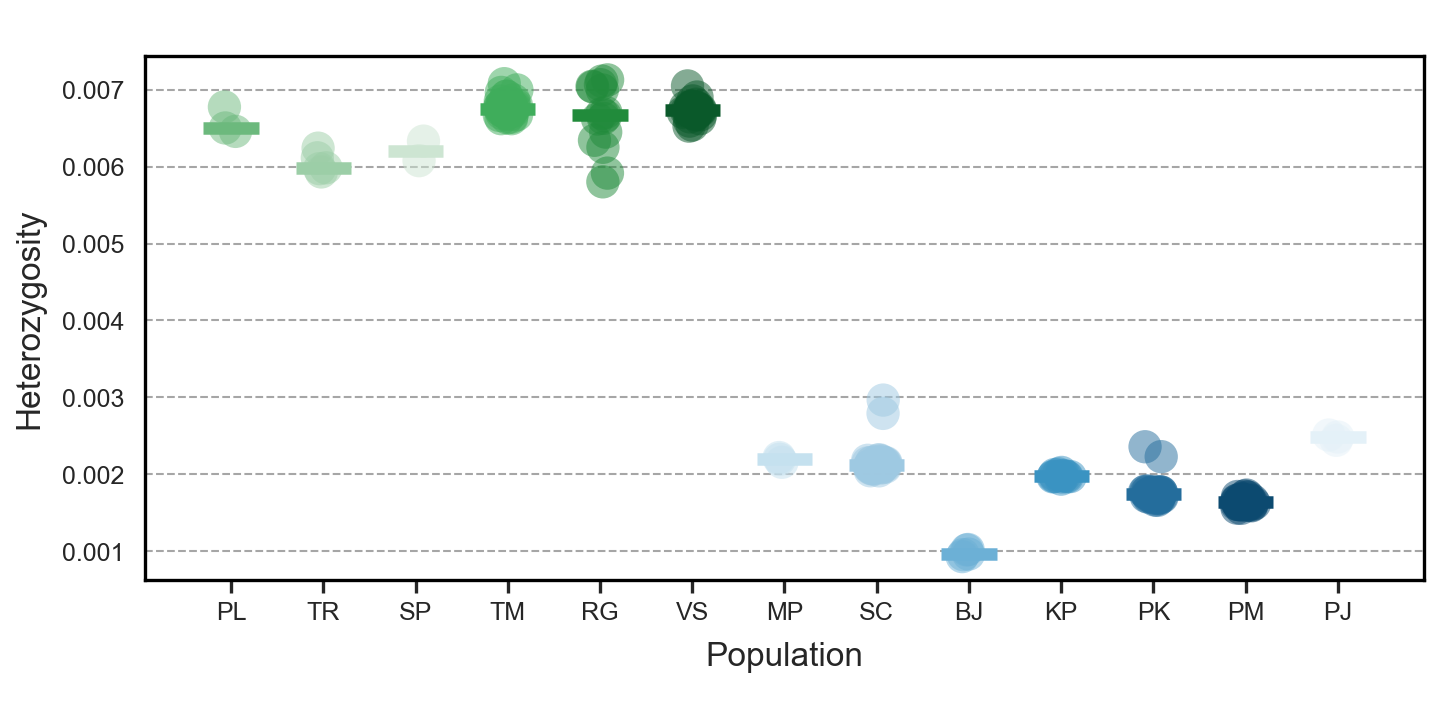

**Figure S2:** Mean genome-wide heterozygosity of *P. siculus* individuals from mainland (green shades) and island (blue shades) populations, estimated with ANGSD v0.941 from genotype likelihoods and folded site frequency spectra computed in 100 kb non-overlapping windows. Points show individual mean heterozygosity (averaged across genome windows). Horizontal lines represent median population-level heterozygosity.

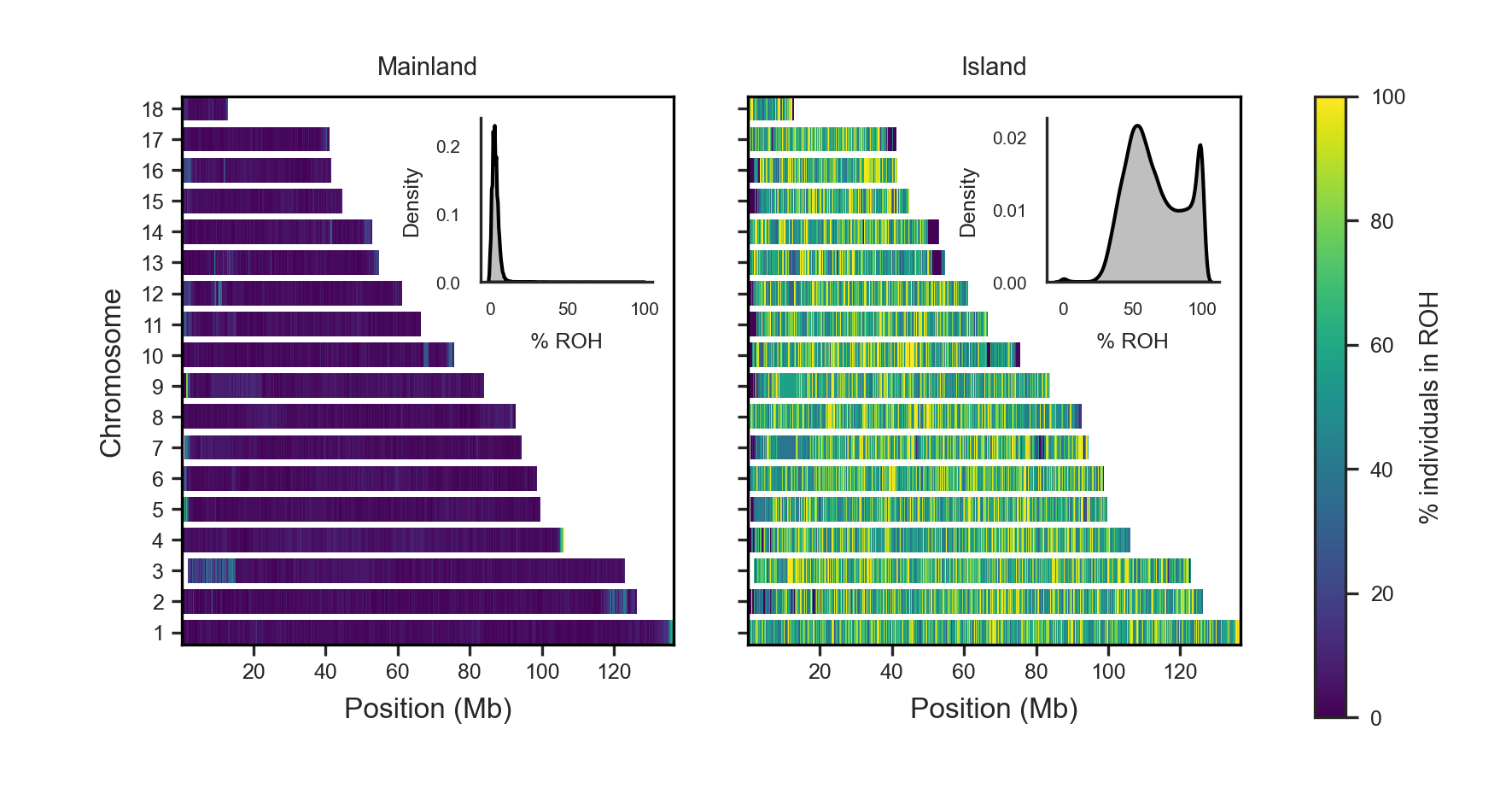

**Figure S3:** Genome-wide distribution of runs of homozygosity (ROH) across mainland and island populations of *P. siculus*. The genome was partitioned into non-overlapping windows of 1500 SNPs, and for each window the proportion of individuals in ROH was calculated for mainland and island groups separately. Colours represent the percentage of individuals with ROH per window, plotted along chromosomes. Density plots summarise the distribution of ROH frequencies across windows.

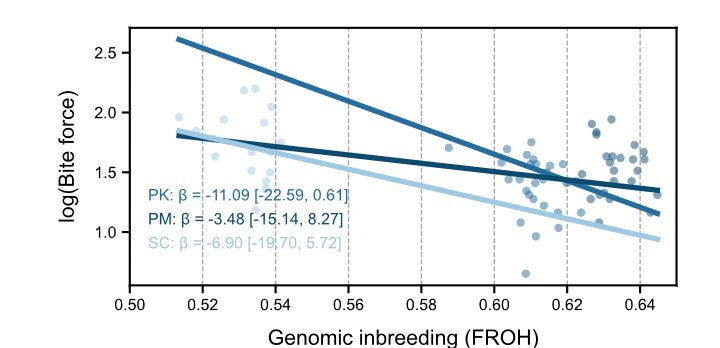

**Figure S4.** Association between genomic inbreeding (F_ROH_) and maximum bite force among island *P. siculus*. Points represent individual lizards, with posterior mean regression lines shown for Pod Kopište (PK), Pod Mrčaru (PM), and Sušac (SC). The estimated slopes (β) and 95% credible intervals are shown for each population. All population-specific slopes were negative, although their 95% credible intervals overlapped zero, and FROH × population interactions were not supported (Table S7C).

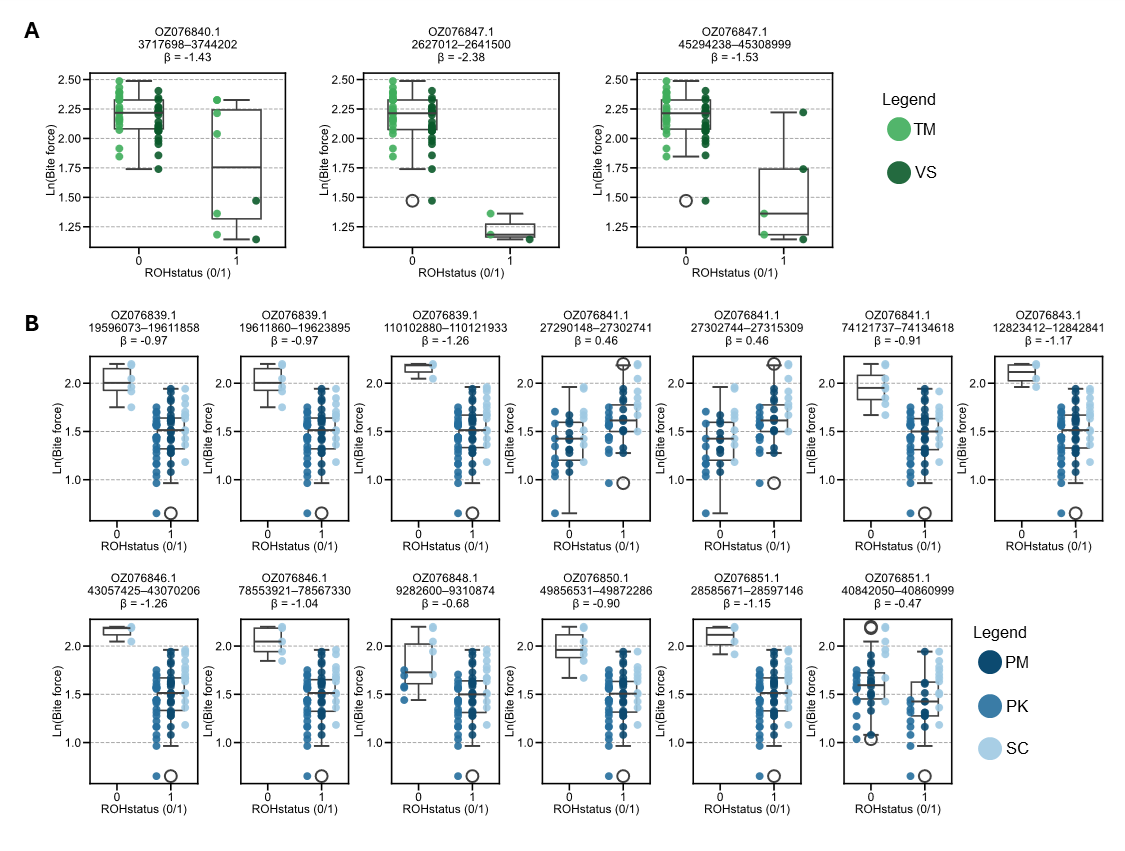

**Figure S5:** Natural log-transformed bite force in relation to ROH status (0=non-ROH, 1=ROH) for genomic windows identified as significant in the ROH-GWAS in male *P. siculus*. Panels show **A.** mainland populations and **B.** island populations. Title of the plots indicate chromosome and genomic coordinates of the significant SNP-window, and the associated effect size. Both boxplots and raw data points are shown with each point representing a male *P. siculus* individual, coloured according to population origin.

**Table S3:** Genetic clusters (K) and their corresponding cross-validation (CV) error based on ADMIXTURE analysis (Alexander et al. 2009). K-values are ordered from lowest to highest.

| **K** | **CV error** |
| --- | --- |
| 2 | 0.10962 |
| 3 | 0.11329 |
| 4 | 0.12162 |
| 8 | 0.12529 |
| 7 | 0.13227 |
| 11 | 0.14642 |
| 10 | 0.14649 |
| 12 | 0.14871 |
| 13 | 0.14982 |
| 14 | 0.16501 |
| 1 | 0.18699 |
| 6 | 0.21176 |
| 9 | 0.2287 |
| 5 | 0.26601 |

**Table S4:** Dsuite results of significant patterns of gene flow (p<0.05) (command: Dtrios, outgroup: *P. muralis*, output: _BBAA.txt, (Malinsky et al. 2021).

| **P1** | **P2** | **P3** | **Dstatistic** | **Z-score** | **p-value** | **f4-ratio** | **BBAA** | **ABBA** | **BABA** |
| --- | --- | --- | --- | --- | --- | --- | --- | --- | --- |
| MP | BJ | PJ | 0.193561 | 13.5709 | 2.3E-16 | 0.294787 | 1232910 | 911551 | 615896 |
| BJ | PJ | SP | 0.100606 | 9.62802 | 2.3E-16 | 0.058669 | 3756820 | 732037 | 598206 |
| BJ | PJ | TR | 0.0835 | 9.23946 | 2.3E-16 | 0.083276 | 3718110 | 718798 | 608010 |
| RG | TR | BJ | 0.032301 | 8.83822 | 2.3E-16 | 0.021144 | 1831220 | 1629000 | 1527060 |
| TM | TR | BJ | 0.033265 | 11.6291 | 2.3E-16 | 0.021773 | 1831640 | 1631490 | 1526450 |
| MP | KP | PJ | 0.189138 | 25.8086 | 2.3E-16 | 0.285886 | 1245040 | 902442 | 615366 |
| KP | PJ | PL | 0.064959 | 11.3879 | 2.3E-16 | 0.052387 | 3760430 | 701611 | 616020 |
| KP | PJ | RG | 0.044962 | 8.41686 | 2.3E-16 | 0.100458 | 3816580 | 673984 | 615985 |
| KP | PJ | SP | 0.095732 | 11.9447 | 2.3E-16 | 0.056176 | 3750130 | 732016 | 604105 |
| KP | PJ | TM | 0.049744 | 10.2887 | 2.3E-16 | 0.150391 | 3817940 | 678531 | 614223 |
| KP | PJ | TR | 0.07891 | 12.8577 | 2.3E-16 | 0.079333 | 3711790 | 719007 | 613833 |
| RG | TR | KP | 0.032405 | 8.79972 | 2.3E-16 | 0.025199 | 1828750 | 1632040 | 1529590 |
| TM | TR | KP | 0.033361 | 11.4983 | 2.3E-16 | 0.025938 | 1829160 | 1634530 | 1528990 |
| MP | PK | PJ | 0.193462 | 22.564 | 2.3E-16 | 0.291642 | 1242920 | 903311 | 610455 |
| MP | PJ | PL | 0.054796 | 13.1827 | 2.3E-16 | 0.049989 | 3554580 | 783764 | 702332 |
| MP | PM | PJ | 0.191178 | 19.1343 | 2.3E-16 | 0.288576 | 1242910 | 902760 | 612983 |
| MP | PJ | RG | 0.034303 | 8.79975 | 2.3E-16 | 0.087836 | 3608110 | 753599 | 703613 |
| MP | SC | PJ | 0.187333 | 28.0591 | 2.3E-16 | 0.284434 | 1242750 | 905136 | 619518 |
| MP | PJ | SP | 0.08394 | 12.5279 | 2.3E-16 | 0.055499 | 3545390 | 815041 | 688809 |
| MP | PJ | TM | 0.038826 | 11.8912 | 2.3E-16 | 0.135006 | 3609650 | 758330 | 701646 |
| MP | PJ | TR | 0.068892 | 15.3689 | 2.3E-16 | 0.078169 | 3507460 | 802689 | 699219 |
| MP | PJ | VS | 0.034625 | 8.24983 | 2.3E-16 | 0.108904 | 3598750 | 754845 | 704321 |
| RG | TR | MP | 0.030378 | 8.49583 | 2.3E-16 | 0.023484 | 1823960 | 1629050 | 1533000 |
| TM | TR | MP | 0.031465 | 11.2677 | 2.3E-16 | 0.024316 | 1824470 | 1631640 | 1532090 |
| PK | PJ | PL | 0.065972 | 11.1392 | 2.3E-16 | 0.052941 | 3763820 | 699213 | 612666 |
| PK | PJ | RG | 0.046121 | 8.30306 | 2.3E-16 | 0.102382 | 3820170 | 671773 | 612540 |
| PK | PJ | SP | 0.096849 | 12.0517 | 2.3E-16 | 0.056566 | 3753550 | 729635 | 600785 |
| PK | PJ | TM | 0.050881 | 10.0558 | 2.3E-16 | 0.152731 | 3821500 | 676299 | 610810 |
| PK | PJ | TR | 0.080066 | 12.8586 | 2.3E-16 | 0.080071 | 3715210 | 716632 | 610383 |
| PM | PJ | PL | 0.06794 | 10.3579 | 2.3E-16 | 0.054529 | 3763310 | 701786 | 612493 |
| RG | PL | PJ | 0.025946 | 11.1058 | 2.3E-16 | 0.021454 | 1800270 | 1636790 | 1554010 |
| SC | PJ | PL | 0.065965 | 12.0627 | 2.3E-16 | 0.053229 | 3760660 | 703278 | 616236 |
| TM | PL | PJ | 0.024878 | 9.83826 | 2.3E-16 | 0.020646 | 1794260 | 1639710 | 1560100 |
| VS | PL | PJ | 0.022659 | 8.99383 | 2.3E-16 | 0.018765 | 1802300 | 1629630 | 1557410 |
| PM | PJ | SP | 0.098863 | 11.5217 | 2.3E-16 | 0.057769 | 3753060 | 732246 | 600488 |
| PM | PJ | TM | 0.053004 | 9.23292 | 2.3E-16 | 0.158339 | 3821010 | 678897 | 610551 |
| PM | PJ | TR | 0.081855 | 11.8912 | 2.3E-16 | 0.081854 | 3714650 | 719162 | 610336 |
| SC | PJ | RG | 0.045524 | 8.92671 | 2.3E-16 | 0.101724 | 3816510 | 675349 | 616537 |
| RG | TR | PJ | 0.046508 | 12.6723 | 2.3E-16 | 0.038748 | 1774030 | 1682670 | 1533110 |
| SC | PJ | SP | 0.096718 | 12.3269 | 2.3E-16 | 0.056792 | 3750350 | 733658 | 604257 |
| SC | PJ | TM | 0.05035 | 11.0308 | 2.3E-16 | 0.152128 | 3817890 | 679918 | 614732 |
| SC | PJ | TR | 0.079933 | 13.8361 | 2.3E-16 | 0.080371 | 3712030 | 720686 | 614001 |
| SC | PJ | VS | 0.045564 | 8.55734 | 2.3E-16 | 0.124747 | 3806900 | 676338 | 617391 |
| TM | TR | PJ | 0.045455 | 17.0495 | 2.3E-16 | 0.037952 | 1772480 | 1683200 | 1536830 |
| VS | TR | PJ | 0.043256 | 10.4629 | 2.3E-16 | 0.036102 | 1775520 | 1675780 | 1536820 |
| RG | TR | PK | 0.03247 | 8.81368 | 2.3E-16 | 0.024383 | 1829160 | 1631370 | 1528760 |
| TM | TR | PK | 0.033409 | 11.5452 | 2.3E-16 | 0.025086 | 1829560 | 1633840 | 1528200 |
| SP | TR | PL | 0.040553 | 9.97649 | 2.3E-16 | 0.108743 | 1820030 | 1708610 | 1575430 |
| RG | TR | PM | 0.032562 | 8.89633 | 2.3E-16 | 0.02396 | 1831650 | 1631270 | 1528380 |
| TM | TR | PM | 0.033501 | 11.6838 | 2.3E-16 | 0.024648 | 1832040 | 1633730 | 1527810 |
| RG | TR | SC | 0.032183 | 8.73685 | 2.3E-16 | 0.026003 | 1829790 | 1631580 | 1529830 |
| TM | TR | SC | 0.033162 | 11.4717 | 2.3E-16 | 0.026788 | 1830220 | 1634080 | 1529180 |
| KP | PJ | VS | 0.045 | 8.02322 | 1.03E-15 | 0.12323 | 3806970 | 674965 | 616834 |
| PK | PJ | VS | 0.046067 | 7.97113 | 1.57E-15 | 0.125304 | 3810500 | 672709 | 613459 |
| RG | SP | PJ | 0.037114 | 7.8912 | 2.99E-15 | 0.031313 | 1749610 | 1686840 | 1566110 |
| PM | PJ | RG | 0.048258 | 7.80148 | 6.12E-15 | 0.106791 | 3819670 | 674367 | 612276 |
| BJ | PJ | PL | 0.069787 | 7.60611 | 2.82E-14 | 0.055863 | 3766840 | 701497 | 609973 |
| VS | SP | PJ | 0.033922 | 7.57055 | 3.72E-14 | 0.028659 | 1751560 | 1679370 | 1569170 |
| PM | PJ | VS | 0.048168 | 7.47829 | 7.53E-14 | 0.130481 | 3809990 | 675284 | 613219 |
| VS | TR | PM | 0.02922 | 7.34238 | 2.1E-13 | 0.021534 | 1833250 | 1624480 | 1532240 |
| VS | TR | BJ | 0.029032 | 7.31352 | 2.6E-13 | 0.019027 | 1832890 | 1622290 | 1530750 |
| VS | TR | PK | 0.029142 | 7.27447 | 3.48E-13 | 0.021919 | 1830780 | 1624590 | 1532590 |
| VS | TR | KP | 0.029116 | 7.25691 | 3.96E-13 | 0.022679 | 1830410 | 1625310 | 1533340 |
| VS | TR | SC | 0.028895 | 7.19347 | 6.32E-13 | 0.023387 | 1831440 | 1624830 | 1533570 |
| TM | PL | BJ | 0.01828 | 7.12476 | 1.04E-12 | 0.011921 | 1847670 | 1601520 | 1544020 |
| RG | PL | BJ | 0.017325 | 7.10245 | 1.23E-12 | 0.011283 | 1851950 | 1596890 | 1542500 |
| TM | SP | PJ | 0.036123 | 7.03892 | 1.94E-12 | 0.030526 | 1751340 | 1686570 | 1568970 |
| TM | PL | PK | 0.018579 | 7.0359 | 1.98E-12 | 0.013899 | 1845430 | 1604120 | 1545600 |
| RG | PL | PK | 0.017648 | 7.02414 | 2.15E-12 | 0.013186 | 1849710 | 1599480 | 1544000 |
| TM | PL | PM | 0.018617 | 7.01558 | 2.29E-12 | 0.013647 | 1847980 | 1603920 | 1545290 |
| RG | PL | PM | 0.017686 | 6.98063 | 2.94E-12 | 0.012948 | 1852260 | 1599280 | 1543690 |
| RG | PL | KP | 0.017551 | 6.98035 | 2.94E-12 | 0.013579 | 1849330 | 1600060 | 1544860 |
| TM | PL | SC | 0.018317 | 6.97374 | 3.09E-12 | 0.014741 | 1846070 | 1604280 | 1546560 |
| TM | PL | KP | 0.018499 | 6.96106 | 3.38E-12 | 0.01433 | 1845060 | 1604720 | 1546430 |
| RG | PL | SC | 0.017346 | 6.93692 | 4.01E-12 | 0.013944 | 1850320 | 1599600 | 1545050 |
| VS | TR | MP | 0.027219 | 6.90235 | 5.12E-12 | 0.021072 | 1825710 | 1622420 | 1536440 |
| PL | TR | PJ | 0.021651 | 6.69659 | 2.13E-11 | 0.017676 | 1963540 | 1574670 | 1507930 |
| TM | PL | MP | 0.017376 | 6.62427 | 3.49E-11 | 0.013384 | 1839800 | 1603740 | 1548960 |
| RG | PL | MP | 0.016296 | 6.58974 | 4.41E-11 | 0.012539 | 1843960 | 1598960 | 1547680 |
| BJ | PJ | TM | 0.054028 | 6.25791 | 3.9E-10 | 0.16078 | 3824080 | 678237 | 608706 |
| VS | PL | BJ | 0.014035 | 5.65168 | 1.59E-08 | 0.009149 | 1854180 | 1589900 | 1545890 |
| VS | PL | PK | 0.0143 | 5.5945 | 2.21E-08 | 0.010699 | 1851860 | 1592410 | 1547510 |
| VS | PL | PM | 0.014324 | 5.57048 | 2.54E-08 | 0.010501 | 1854400 | 1592200 | 1547230 |
| VS | PL | KP | 0.014242 | 5.54635 | 2.92E-08 | 0.011035 | 1851500 | 1593020 | 1548280 |
| VS | PL | SC | 0.014038 | 5.4995 | 3.81E-08 | 0.011302 | 1852480 | 1592540 | 1548450 |
| BJ | PJ | RG | 0.049214 | 5.49449 | 3.92E-08 | 0.10857 | 3822720 | 673678 | 610479 |
| RG | VS | BJ | 0.003381 | 5.43787 | 5.39E-08 | 0.002159 | 1897090 | 1545040 | 1534630 |
| BJ | PJ | VS | 0.049314 | 5.40153 | 6.61E-08 | 0.133054 | 3813110 | 674669 | 611255 |
| RG | VS | PM | 0.003455 | 5.25271 | 1.5E-07 | 0.00248 | 1897760 | 1546750 | 1536100 |
| RG | VS | KP | 0.003401 | 5.21981 | 1.79E-07 | 0.00258 | 1894710 | 1547650 | 1537160 |
| RG | VS | PK | 0.00344 | 5.20677 | 1.92E-07 | 0.00252 | 1895190 | 1547000 | 1536390 |
| SP | PL | RG | 0.019344 | 5.17678 | 2.26E-07 | 0.187036 | 1810090 | 1651330 | 1588660 |
| VS | PL | MP | 0.013119 | 5.13969 | 2.75E-07 | 0.010107 | 1846250 | 1592040 | 1550810 |
| RG | VS | SC | 0.0034 | 5.1346 | 2.83E-07 | 0.00268 | 1895470 | 1547600 | 1537120 |
| RG | VS | MP | 0.003266 | 5.06111 | 4.17E-07 | 0.002465 | 1888100 | 1548840 | 1538760 |
| PL | TR | BJ | 0.015758 | 5.0302 | 4.9E-07 | 0.009972 | 2031420 | 1531770 | 1484250 |
| SP | PL | VS | 0.019247 | 5.0067 | 5.54E-07 | 0.270476 | 1803220 | 1653650 | 1591200 |
| PL | TR | PM | 0.015655 | 4.96488 | 6.87E-07 | 0.011156 | 2031560 | 1533470 | 1486200 |
| PL | TR | PK | 0.015598 | 4.88811 | 1.02E-06 | 0.011347 | 2029100 | 1533600 | 1486490 |
| PL | TR | SC | 0.015614 | 4.87003 | 1.12E-06 | 0.01223 | 2030040 | 1534120 | 1486950 |
| PL | TR | KP | 0.015632 | 4.86331 | 1.15E-06 | 0.01178 | 2028750 | 1534340 | 1487110 |
| PL | TR | MP | 0.014815 | 4.63049 | 3.65E-06 | 0.011082 | 2025210 | 1532740 | 1487990 |
| SP | PL | TM | 0.015962 | 4.37319 | 1.22E-05 | 0.28627 | 1814060 | 1646370 | 1594640 |
| SP | TR | TM | 0.017223 | 4.33381 | 1.47E-05 | 0.294315 | 1981730 | 1570760 | 1517570 |
| RG | VS | PJ | 0.003399 | 4.26579 | 1.99E-05 | 0.002746 | 1853110 | 1564450 | 1553850 |
| RG | VS | TR | 0.00255 | 4.1226 | 3.75E-05 | 0.007426 | 1699980 | 1650010 | 1641620 |
| SP | TR | MP | 0.01735 | 3.90293 | 9.5E-05 | 0.012721 | 2111000 | 1507400 | 1455980 |
| SP | TR | RG | 0.018564 | 3.76754 | 0.000165 | 0.170933 | 1980550 | 1571660 | 1514370 |
| SP | TR | PM | 0.017392 | 3.76748 | 0.000165 | 0.01216 | 2116750 | 1507450 | 1455910 |
| SP | TR | PK | 0.01728 | 3.76691 | 0.000165 | 0.012333 | 2114300 | 1507580 | 1456370 |
| SP | TR | KP | 0.017318 | 3.74285 | 0.000182 | 0.012804 | 2113960 | 1508330 | 1456980 |
| SP | TR | BJ | 0.017479 | 3.71229 | 0.000205 | 0.010854 | 2116670 | 1505900 | 1454160 |
| SP | TR | SC | 0.017314 | 3.69042 | 0.000224 | 0.013305 | 2115280 | 1508150 | 1456820 |
| TM | SP | PK | 0.017004 | 3.67365 | 0.000239 | 0.012888 | 1815210 | 1621440 | 1567220 |
| TM | SP | PM | 0.016994 | 3.65413 | 0.000258 | 0.01262 | 1817760 | 1621080 | 1566910 |
| TM | SP | KP | 0.016919 | 3.64394 | 0.000268 | 0.013279 | 1814860 | 1622050 | 1568070 |
| TM | SP | SC | 0.016728 | 3.58629 | 0.000335 | 0.01364 | 1815910 | 1621620 | 1568260 |
| RG | SP | PK | 0.016039 | 3.58253 | 0.00034 | 0.012164 | 1811540 | 1619780 | 1568640 |
| RG | SP | PM | 0.016028 | 3.57273 | 0.000353 | 0.011911 | 1814100 | 1619440 | 1568340 |
| RG | SP | KP | 0.015938 | 3.55074 | 0.000384 | 0.012518 | 1811170 | 1620360 | 1569520 |
| RG | SP | SC | 0.015723 | 3.47958 | 0.000502 | 0.012831 | 1812200 | 1619920 | 1569760 |
| TM | SP | BJ | 0.016677 | 3.46213 | 0.000536 | 0.011018 | 1817540 | 1618820 | 1565710 |
| TM | SP | MP | 0.015028 | 3.4304 | 0.000603 | 0.011723 | 1810280 | 1619210 | 1571260 |
| SP | TR | VS | 0.018199 | 3.41635 | 0.000635 | 0.243553 | 1973820 | 1573330 | 1517080 |
| RG | VS | PL | 0.002793 | 3.40414 | 0.000664 | 0.006649 | 1698160 | 1654440 | 1645220 |
| RG | SP | BJ | 0.01569 | 3.3755 | 0.000737 | 0.010372 | 1813870 | 1617160 | 1567200 |
| RG | SP | MP | 0.013919 | 3.31003 | 0.000933 | 0.010867 | 1806490 | 1617420 | 1573010 |
| KP | PM | BJ | 0.015603 | 3.23285 | 0.001226 | 0.023458 | 630561 | 630428 | 611058 |
| VS | SP | PK | 0.012746 | 2.95462 | 0.003131 | 0.00968 | 1813750 | 1612560 | 1571970 |
| VS | SP | PM | 0.012722 | 2.94399 | 0.00324 | 0.009466 | 1816300 | 1612200 | 1571700 |
| VS | SP | KP | 0.012683 | 2.92715 | 0.003421 | 0.009974 | 1813410 | 1613170 | 1572760 |
| VS | SP | SC | 0.012471 | 2.86128 | 0.004219 | 0.010191 | 1814440 | 1612720 | 1572990 |
| SC | MP | RG | 0.008442 | 2.85945 | 0.004244 | 0.015175 | 4054240 | 524119 | 515344 |
| RG | VS | SP | 0.00289 | 2.82642 | 0.004707 | 0.004517 | 1743060 | 1636960 | 1627530 |
| VS | SP | BJ | 0.012454 | 2.7618 | 0.005748 | 0.008241 | 1816120 | 1609970 | 1570370 |
| KP | PK | SC | 0.005511 | 2.73874 | 0.006167 | 0.378225 | 729772 | 616394 | 609637 |
| SC | MP | VS | 0.008051 | 2.7356 | 0.006227 | 0.017724 | 4044470 | 524432 | 516055 |
| SC | MP | TM | 0.008128 | 2.73294 | 0.006277 | 0.019712 | 4057490 | 523877 | 515429 |
| KP | PK | BJ | 0.011546 | 2.7059 | 0.006812 | 0.017347 | 633087 | 627440 | 613116 |
| VS | SP | MP | 0.010794 | 2.64936 | 0.008064 | 0.008438 | 1808860 | 1610370 | 1575970 |
| TM | VS | PL | 0.005468 | 2.60031 | 0.009314 | 0.013007 | 1683580 | 1668110 | 1649960 |
| PL | SP | PJ | 0.012021 | 2.45222 | 0.014198 | 0.010082 | 1884770 | 1600750 | 1562720 |
| PM | MP | RG | 0.011696 | 2.31709 | 0.020499 | 0.020771 | 4059290 | 522457 | 510378 |
| TM | VS | SC | 0.004398 | 2.31261 | 0.020744 | 0.003484 | 1877530 | 1557920 | 1544270 |
| TM | VS | PK | 0.004399 | 2.30676 | 0.021068 | 0.003239 | 1877200 | 1557270 | 1543630 |
| TM | VS | PM | 0.004413 | 2.30574 | 0.021125 | 0.003184 | 1879760 | 1557010 | 1543330 |
| TM | VS | MP | 0.004372 | 2.30527 | 0.021152 | 0.003316 | 1870270 | 1559250 | 1545680 |
| KP | MP | RG | 0.007702 | 2.30322 | 0.021267 | 0.013801 | 4057960 | 521369 | 513399 |
| TM | VS | KP | 0.004377 | 2.29996 | 0.02145 | 0.003337 | 1876750 | 1557940 | 1544360 |
| TM | VS | BJ | 0.004364 | 2.28534 | 0.022293 | 0.002801 | 1879130 | 1555300 | 1541790 |
| PM | MP | TM | 0.011263 | 2.22531 | 0.02606 | 0.026941 | 4062480 | 522154 | 510523 |
| TM | VS | TR | 0.003158 | 2.2178 | 0.026568 | 0.009237 | 1683070 | 1661350 | 1650890 |
| PM | MP | VS | 0.011143 | 2.21747 | 0.026591 | 0.024214 | 4049440 | 522695 | 511174 |
| PK | MP | RG | 0.008941 | 2.21569 | 0.026713 | 0.015927 | 4059410 | 520043 | 510826 |
| KP | MP | VS | 0.007307 | 2.21292 | 0.026903 | 0.016042 | 4048210 | 521703 | 514134 |
| KP | MP | TM | 0.007321 | 2.16699 | 0.030236 | 0.017713 | 4061180 | 521100 | 513525 |
| PK | MP | VS | 0.008432 | 2.09943 | 0.035779 | 0.018396 | 4049600 | 520310 | 511609 |
| PK | MP | TM | 0.008506 | 2.09738 | 0.03596 | 0.020443 | 4062610 | 519750 | 510982 |
| BJ | PK | SC | 0.01173 | 1.99718 | 0.045805 | 0.566323 | 730141 | 622476 | 608043 |
| TM | RG | PL | 0.00269 | 1.98146 | 0.04754 | 0.0064 | 1684890 | 1663590 | 1654660 |

**Table S5:** Population-level inbreeding estimates based on individual inbreeding coefficients (F) calculated from deviations from Hardy-Weinberg equilibrium using the --het function in VCFtools (Danecek et al., 2011). Values represent the mean (± SD) of individual estimates per population.

| **Population** | **F (± SD)** |
| --- | --- |
| BJ | -0.024 (± 0.043) |
| KP | 0.0036 (± 0.013) |
| MP | 0.0005 (± 0.022) |
| PJ | 0.0036 (± 0.0065) |
| PK | -0.010 (± 0.11) |
| PL | 0.012 (± 0.020) |
| PM | -0.00092 (± 0.030) |
| RG | 0.038 (± 0.043) |
| SC | -0.015 (± 0.11) |
| SP | 0.0036 (± 0.0023) |
| TM | 0.015 (± 0.013) |
| TR | 0.0095 (± 0.018) |
| VS | 0.013 (± 0.010) |

**Table S6:** Population-average F_ROH_, morphometrics, and performance (PK, PM, SC = island; TM, VS = mainland). See Supplementary Methods for details on how the performance traits were measured. SVL = snout-vent-length.

| **Pop** | **F_ROH_** | **SVL (mm)** | **Head size (mm)** | **Ticks (N°)** | **Mites**  **(N°)** | **Bite force (N)** | **Sprint speed (cm/s)** | **Sperm length (µm)** | **Sperm velocity (µm/s)** | **Sperm count (#)** | **PHA swelling (mm)** | **# Blood parasites** | **# Trials to spatial learning** |
| --- | --- | --- | --- | --- | --- | --- | --- | --- | --- | --- | --- | --- | --- |
| PK | 0.61  ± 0.0071 | 61.67  ± 2.47 | 10.37  ± 0.35 | 0 | 0 | 4.12  ± 0.99 | 160.78  ± 36.97 | 139.72  ± 4.11 | 53.57  ± 7.04 | 7.21  ± 10.49 | 0.87  ± 0.24 | 1.33  ± 3.37 | 13.08  ± 5.16 |
| PM | 0.63  ± 0.0069 | 65.69  ± 2.45 | 11.18  ± 0.29 | 0 | 0 | 4.7  ± 1.05 | 165.16  ± 31.70 | 150.92  ± 11.51 | 48.01  ± 9.32 | 7.07  ± 6.53 | 0.86  ± 0.27 | 0.19  ± 0.40 | 9.63  ± 4.30 |
| SC | 0.53  ± 0.0081 | 64.68  ± 2.76 | 10.88  ± 0.64 | 0.04  ± 0.20 | 0 | 5.9  ± 1.63 | 183.42  ± 38.63 | 145.11  ± 20.98 | 55.84  ± 7.99 | 6.32  ± 7.33 | 0.83  ± 0.22 | 0.39  ± 0.89 | 15.92  ± 3.83 |
| TM | 0.09  ± 0.0064 | 71.21  ± 5.01 | 11.93  ± 0.84 | 0 | 0.36  ± 1.89 | 9.07  ± 2.05 | 248.31  ± 49.04 | 153.38  ± 2.94 | 51.75  ± 5.38 | 15.71  ± 13.29 | 0.98  ± 0.30 | 0.04  ± 0.19 | 13.93  ± 5.47 |
| VS | 0.09  ± 0.0062 | 70.16  ± 3.97 | 11.51  ± 0.71 | 0 | 10.19  ± 17.12 | 8.16  ± 1.80 | 240.16  ± 37.03 | 135.50  ± 12.06 | 50.35  ± 5.93 | 19.56  ± 14.75 | 0.93  ± 0.13 | 0.88  ± 1.70 | 12.81  ± 6.13 |

**Table S7: A.** MCMCglmm results for the effect of F_ROH_ on lizard bite force (BF, log-transformed) for mainland and island lizards with head size (log-transformed) as a covariate, fitted with a gaussian distribution (GRM = genetic relatedness matrix). The island slope is given by the F_ROH_ main effect, while the mainland slope was calculated by adding the F_ROH_ main effect and the F_ROH_ × type interaction using posterior samples. **B.** MCMCglmm results for the within-population association between F_ROH_ and bite force among island lizards, with F_ROH_ centred around the population mean. **C.** MCMCglmm results for the association between F_ROH_ and bite force among island lizards, including FROH × population interactions

| A. **BF ~ F_ROH_** Non-informative priors | | | |
| --- | --- | --- | --- |
|  | Effective N | Posterior mean [95% CI] | pMCMC |
| Random effect |  |  |  |
| GRM | 8039 | 0.011 [0.00016, 0.041] | - |
| Population | 9990 | 0.014 [0.00015, 0.042] | - |
| Units | 10880 | 0.034 [0.026, 0.044] | - |
| Fixed effect |  |  |  |
| Intercept | 11649 | 0.18 [-1.44, 1.74] | 0.87 |
| **F_ROH_** | **9990** | **-3.55 [-6.20, -1.12]** | **0.0030** |
| **Head size** | **10158** | **0.32 [0.27, 0.38]** | **< 0.0001** |
| **Origin-mainland** | **9990** | **-2.21 [-3.99, -0.63]** | **0.0056** |
| F_ROH_:Origin-mainland | 9990 | 7.93 [-0.24, 16.98] | 0.071 |

| B. **Island-only BF ~ F_ROH-centred_** Non-informative priors | | | |
| --- | --- | --- | --- |
|  | Effective N | Posterior mean [95% CI] | pMCMC |
| Random effect |  |  |  |
| GRM | 7000 | 0.14 [0.00016, 0.69] | - |
| Units | 8080 | 0.038 [0.011, 0.061] | - |
| Fixed effect |  |  |  |
| **Intercept** | **10240** | **-2.34 [-3.69, 1.03]** | **0.0010** |
| **F_ROH-centred_** | **9990** | **-7.12 [-14.14, -0.35]** | **0.042** |
| Population-PM | 9682 | -0.14 [-0.30, 0.022] | 0.083 |
| **Population-SC** | **9548** | **0.18 [0.026, 0.34]** | **0.025** |
| **Head size** | **10281** | **0.36 [0.23, 0.49]** | **< 0.0001** |

| C. **Island-only BF ~ F_ROH_** Non-informative priors | | | |
| --- | --- | --- | --- |
|  | Effective N | Posterior mean [95% CI] | pMCMC |
| Random effect |  |  |  |
| GRM | 6973 | 0.16 [0.00024, 0.72] | - |
| Units | 7796 | 0.039 [0.0094, 0.061] | - |
| Fixed effect |  |  |  |
| Intercept | 9990 | 4.46 [-2.80, 11.84] | 0.23 |
| F_ROH_ | 9990 | -11.08 [-22.81, 0.81] | 0.067 |
| Population-PM | 9990 | -4.69 [-15.06, 5.53] | 0.37 |
| Population-SC | 9706 | -2.95 [-13.21, 6.73] | 0.56 |
| **Head size** | **9673** | **0.35 [0.22, 0.48]** | **< 0.0001** |
| F_ROH_:Population-PM | 9990 | 7.58 [-9.06, 24.06] | 0.37 |
| F_ROH_:Population-SC | 9686 | 4.28 [-13.21, 21.79] | 0.63 |

**Table S8:** MCMCglmm results for the effect of F_ROH_ on lizard sprint speed (log-transformed) for mainland and island lizards with SVL (log-transformed) as a covariate, fitted with a gaussian distribution (GRM = genetic relatedness matrix).

| **Sprint ~ F_ROH_** Non-informative priors | | | |
| --- | --- | --- | --- |
|  | Effective N | Posterior mean [95% CI] | pMCMC |
| Random effect |  |  |  |
| GRM | 7497 | 0.019 [0.00016, 0.074] | - |
| Population | 9990 | 0.018 [0.00017, 0.063] | - |
| Units | 8833 | 0.042 [0.031, 0.054] | - |
| Fixed effect |  |  |  |
| **Intercept** | **9990** | **5.41 [2.00, 8.74]** | **0.0056** |
| F_ROH_ | 9990 | -0.40 [-3.05, 2.90] | 0.60 |
| SVL | 9990 | -0.015 [-0.68, 0.66] | 0.97 |
| Origin-mainland | 9990 | -0.028 [-1.94, 2.00] | 0.87 |
| F_ROH_:Origin-mainland | 9990 | 2.22 [-7.34, 11.80] | 0.65 |

**Table S9:** MCMCglmm results for the effect of F_ROH_ on lizard sperm count for mainland and island lizards with SVL (log-transformed) as a covariate, fitted with a Poisson distribution (GRM = genetic relatedness matrix).

| **Sperm count ~ F_ROH_** Non-informative priors | | | |
| --- | --- | --- | --- |
|  | Effective N | Posterior mean [95% CI] | pMCMC |
| Random effect |  |  |  |
| GRM | 5352 | 0.23 [0.00023, 1.18] | - |
| Population | 9990 | 0.14 [0.00016, 0.36] | - |
| Units | 9403 | 0.75 [0.51, 1.03] | - |
| Fixed effect |  |  |  |
| Intercept | 9990 | -1.60 [-17.12, 13.01] | 0.84 |
| F_ROH_ | 8444 | 3.66 [-5.51, 13.46] | 0.39 |
| SVL | 10705 | 0.22 [-3.06, 3.62] | 0.90 |
| Origin-mainland | 9990 | 4.23 [-2.51, 11.29] | 0.19 |
| F_ROH_:Origin-mainland | 9990 | -14.76 [-57.01, 26.77] | 0.49 |

**Table S10:** MCMCglmm results for the effect of F_ROH_ on lizard sperm size (total sperm length) for mainland and island lizards with SVL (log-transformed) and sperm count as covariates, fitted with a gaussian distribution (GRM = genetic relatedness matrix).

| **Sperm size ~ F_ROH_** Non-informative priors | | | |
| --- | --- | --- | --- |
|  | Effective N | Posterior mean [95% CI] | pMCMC |
| Random effect |  |  |  |
| GRM | 3622 | 10.70 [0.00015, 61.61] | - |
| Population | 9990 | 439.7 [6.29, 1361] | - |
| Units | 8766 | 123.8 [92.04, 159.9] | - |
| Fixed effect |  |  |  |
| Intercept | 9093 | 177.49 [-95.03, 447.02] | 0.18 |
| F_ROH_ | 9990 | -142.01 [-484.99, 137.23] | 0.36 |
| SVL | 9179 | 12.80 [-30.55, 54.16] | 0.55 |
| Sperm count | 9990 | -0.078 [-0.27, 0.11] | 0.41 |
| Origin-mainland | 9990 | -120.00 [-330.25, 61.99] | 0.19 |
| F_ROH_:Origin-mainland | 9990 | 510.79 [-129.92, 1112.38] | 0.10 |

**Table S11:** MCMCglmm results for the effect of F_ROH_ on lizard sperm velocity (log-transformed) for mainland and island lizards with SVL (log-transformed) and total sperm length as covariates, fitted with a gaussian distribution (GRM = genetic relatedness matrix).

| **Sperm velocity ~ F_ROH_** Non-informative priors | | | |
| --- | --- | --- | --- |
|  | Effective N | Posterior mean [95% CI] | pMCMC |
| Random effect |  |  |  |
| GRM | 9176 | 0.0067 [0.00020, 0.023] | - |
| Population | 9990 | 0.024 [0.00015, 0.061] | - |
| Units | 9990 | 0.018 [0.013, 0.024] | - |
| Fixed effect |  |  |  |
| **Intercept** | **10578** | **3.34 [0.66, 6.17]** | **0.017** |
| F_ROH_ | 9990 | -2.04 [-5.03, 0.45] | 0.067 |
| SVL | 10312 | 0.42 [-0.092, 0.96] | 0.11 |
| Sperm size | 9990 | 0.00043 [-0.0018, 0.0027] | 0.71 |
| **Origin-mainland** | **9990** | **-1.59 [-3.45, 0.053]** | **0.037** |
| F_ROH_:Origin-mainland | 9990 | 5.56 [-1.35, 12.62] | 0.12 |

**Table S12:** MCMCglmm results for the effect of F_ROH_ on PHA immune response for mainland and island lizards with SVL (log-transformed) and haemoparasite count as covariates, fitted with a gaussian distribution (GRM = genetic relatedness matrix). Note: only one individual had a tick infestation, and only individuals from the VS population were infested with mites, as such the prevalence of these ectoparasites was omitted in the model to aid convergence.

| **PHA response ~ F_ROH_** Non-informative priors | | | |
| --- | --- | --- | --- |
|  | Effective N | Posterior mean [95% CI] | pMCMC |
| Random effect |  |  |  |
| GRM | 7778 | 0.023 [0.00023, 0.091] | - |
| Population | 9990 | 0.024 [0.00013, 0.073] | - |
| Units | 9990 | 0.050 [0.037, 0.064] | - |
| Fixed effect |  |  |  |
| Intercept | 9990 | -3.23 [-7.28, 0.46] | 0.081 |
| F_ROH_ | 9707 | 0.93 [-2.04, 4.63] | 0.56 |
| **SVL** | **9990** | **0.85 [0.12, 1.62]** | **0.023** |
| Haemoparasite count | 9990 | -0.0016 [-0.024, 0.021] | 0.89 |
| Origin-mainland | 9650 | 0.21 [-1.86, 2.54] | 0.95 |
| F_ROH_:Origin-mainland | 9990 | 2.86 [-7.88, 14.14] | 0.60 |

**Table S13:** MCMCglmm results for the effect of F_ROH_ on number of trials to successfully reach the learning criterion of spatial learning for mainland and island lizards with SVL (log-transformed) as a covariate, fitted with a Poisson distribution (GRM = genetic relatedness matrix).

| **Spatial learning ~ F_ROH_** Non-informative priors | | | |
| --- | --- | --- | --- |
|  | Effective N | Posterior mean [95% CI] | pMCMC |
| Random effect |  |  |  |
| GRM | 4092 | 0.70 [0.00027, 1.25] | - |
| Population | 9990 | 0.051 [0.00015, 0.18] | - |
| Units | 3364 | 0.024 [0.00019, 0.083] | - |
| Fixed effect |  |  |  |
| Intercept | 9961 | 5.37 [-1.52, 12.35] | 0.14 |
| F_ROH_ | 9990 | -3.28 [-8.72, 2.18] | 0.20 |
| SVL | 10315 | -0.22 [-1.57, 1.21] | 0.75 |
| Origin-mainland | 8920 | -2.25 [-6.00, 1.61] | 0.21 |
| F_ROH_:Origin-mainland | 9632 | 7.21 [-13.97, 28.42] | 0.50 |

**Table S14:** SNP windows in ROHs significantly associated with bite force in mainland and island *P. siculus* from the ROH-GWAS, including overlapping protein-coding genes identified from the *P. muralis* to *P. siculus* Liftoff annotation.

| **Chrom:SNPwindow_start-SNPwindow_end** | **Gene ID** | **Gene biotype** | **Gene coordinates_start** | **Gene coordinates_end** |
| --- | --- | --- | --- | --- |
| ***Mainland*** | | | | |
| OZ076840.1:3717698-3744202 | MEIS1 | Protein coding | 3620685 | 3771217 |
| OZ076847.1:2627012-2641500 | LOC114604855 | Protein coding | 2493668 | 2812430 |
| OZ076847.1:45294238-45308999 | NUP107 | Protein coding | 45280971 | 45305896 |
|  | RAP1B | Protein coding | 45306626 | 45333645 |
| ***Island*** | | | | |
| OZ076839.1:19596073-19611858 | LOC114588424 | Protein coding | 19595092 | 19607591 |
| OZ076839.1:19611860-19623895 | LOC114588406 | Protein coding | 19615790 | 19623401 |
| OZ076839.1:110102880-110121933 | SCN8A | Protein coding | 110025277 | 110135585 |
| OZ076841.1:27290148-27302741 | FRY | Protein coding | 27202783 | 27390649 |
| OZ076841.1:27302744-27315309 | FRY | Protein coding | 27202783 | 27390649 |
| OZ076841.1:74121737-74134618 | NA | NA | NA | NA |
| OZ076843.1:12823412-12842841 | BRINP3 | Protein coding | 12825389 | 13088350 |
| OZ076846.1:43057425-43070206 | NA | NA | NA | NA |
| OZ076846.1:78553921-78567330 | SLC4A11 | Protein coding | 78441475 | 78645013 |
| OZ076848.1:9282600-9310874 | LOC114606797 | Protein coding | 9276644 | 9350771 |
| OZ076850.1:49856531-49872286 | INO80C | Protein coding | 49851328 | 49879540 |
| OZ076851.1:28585671-28597146 | IGDCC3 | Protein coding | 28506402 | 28617167 |
| OZ076851.1:40842050-40860999 | NA | NA | NA | NA |
